# Epigenetic clocks predict prevalence and incidence of leading causes of death and disease burden

**DOI:** 10.1101/2020.01.31.928648

**Authors:** Robert F. Hillary, Anna J. Stevenson, Daniel L. McCartney, Archie Campbell, Rosie M. Walker, David M. Howard, Craig W. Ritchie, Steve Horvath, Caroline Hayward, Andrew M. McIntosh, David J. Porteous, Ian J. Deary, Kathryn L. Evans, Riccardo E. Marioni

## Abstract

Individuals of the same chronological age display different rates of biological ageing. A number of measures of biological age have been proposed which harness age-related changes in DNA methylation profiles. These include methylation-based predictors of chronological age (HorvathAge, HannumAge), all-cause mortality (DNAm PhenoAge, DNAm GrimAge) and telomere length (DNAm Telomere Length). In this study, we test the association between these epigenetic markers of ageing and the prevalence and incidence of the leading causes of disease burden and mortality in high-income countries. Furthermore, we test the clocks’ relationships with phenotypic measures associated with these conditions, including spirometric and biochemical traits. We carry out these analyses in 9,537 individuals from the Generation Scotland: Scottish Family Health Study. We find that DNAm GrimAge outperforms other epigenetic clocks in its associations with self-report disease prevalence and related clinical traits. DNAm GrimAge associates with chronic obstructive pulmonary disease (COPD) prevalence (Odds Ratio = 3.29, P = 3.0 × 10^-4^) and pulmonary spirometry tests (β = [-0.10 to −0.15], P = [1.4 × 10^-4^ to 1.4 × 10^-6^]) at study baseline after adjusting for possibly confounding risk factors including alcohol, body mass index, deprivation, education and smoking. After adjusting for these confounding risk factors, DNAm GrimAge, DNAm PhenoAge and DNAm Telomere Length, measured at study baseline, predict incidence of ICD-10-coded disease states including COPD, type 2 diabetes and cardiovascular disease after thirteen years of follow-up (Hazard Ratios = [0.80 (telomere length) to 2.19 (GrimAge)], P = [9.9 × 10^-4^, 1.9 × 10^-14^]). Our data show that despite accounting for several possible confounding variables, epigenetic markers of ageing predict incidence of common disease. This may have significant implications for their potential utility in clinical settings to complement gold-standard methods of clinical assessment and management.

## Introduction

The sustained increase in global life expectancy and population size has prompted a concomitant elevation in the prevalence of chronic disease states (1). The World Health Organisation specifies ten leading causes of mortality and ten leading causes of disease burden. In high-income countries, six diseases are present in both sets: ischemic heart disease, stroke, lung cancer, Alzheimer’s disease (AD) and other dementias, diabetes and chronic obstructive pulmonary disease (COPD). The remaining four leading causes of mortality are lower respiratory tract diseases, bowel cancer, kidney disease and breast cancer (2). The additional four causes of disease burden are back or neck pain, skin disease, sense organ disease and depression (3). All of these disease states encompass heterogeneous, complex aetiologies resulting in a paucity of effective treatment paradigms. Given the number of individuals affected by such disorders and the associated burden, there is an urgent need for effective molecular predictors in clinical settings that can discriminate individuals on trajectories towards disease.

Ageing is a major risk factor for many common disease states. However, individuals of the same chronological age exhibit disparate rates of biological ageing and susceptibilities to common morbidities and mortality. Differential patterns of biological ageing among individuals may be exploited to identify novel predictors of disease (4). Recently, a number of strategies have been proposed to estimate biological age by leveraging inter-individual variation in DNA methylation (DNAm) profiles. These so-called ‘epigenetic clocks’ correlate strongly with chronological age (5). Moreover, for a given chronological age, an accelerated epigenetic age is associated with an increased risk of mortality and shows cross-sectional relationships with age-related morbidities (6–10).

In this paper, we focus on five epigenetic predictors of ageing. In 2013, Horvath developed a pan-tissue epigenetic clock, termed ‘HorvathAge’, derived from the linear combination of 353 CpG sites in multiple tissues (11). Hannum created a DNAm-based clock termed ‘HannumAge’ based on 71 CpGs in blood tissue (12). Levine *et al.* proposed a predictor of lifespan and health by developing a methylation-based predictor of an individual’s ‘phenotypic age’ (‘DNAm PhenoAge’). Phenotypic age is informed by chronological age as well as haematological and biochemical measures, including creatinine levels and lymphocyte percent (13). Lu *et al.* proposed DNAm GrimAge as a predictor of mortality and demonstrated that it outperforms existing clocks in predicting death and age-related conditions, including cardiovascular disease (14). Furthermore, telomere length is associated with cardiovascular disease, cancer risk and all-cause mortality (15–17). Lu *et al.* proposed a DNAm-based estimator of telomere length termed ‘DNAm Telomere Length’ (DNAm TL) which exhibits stronger associations with lifespan, smoking history and body mass index when compared to phenotypic telomere length as measured by quantitative polymerase chain reaction or Southern blotting (18).

The difference between an individual’s methylation-based age and their chronological age provides a measure of accelerated or decelerated ageing. Higher values of age-adjusted HorvathAge, HannumAge, DNAm PhenoAge and DNAm GrimAge are hypothesised to associate with poorer health outcomes as these clocks were trained to predict lifespan mortality. Lower values of age-adjusted DNAm TL are hypothesised to correlate with poorer health as this reflects shorter telomere length. To date, a number of studies have demonstrated associations between epigenetic clocks and risk of mortality and disease states (19–22), or have provided comparisons of clocks (e.g. HorvathAge vs. HannumAge) (23–27). However, no study has compared all major epigenetic clocks with respect to their association with a broad range of common health conditions.

In this study, we test the association between all five epigenetic clocks and the prevalence, and incidence, of the ten leading causes of mortality and disease burden (as indexed by disability-adjusted life years; DALYs) (2, 3). In addition, we examine their association with continuous traits underlying these conditions, such as lung function tests for Chronic Obstructive Pulmonary Disease (COPD). We utilise DNA methylation array data and electronic health record data from a Scottish cohort: Generation Scotland: Scottish Family Health Study (GS:SFHS or GS). GS is a family-based cohort consisting of over 20,000 individuals with rich health and lifestyle information. Genome-wide methylation data were generated on approximately 10,000 participants making it one of the largest DNAm resources in the world. We examine associations between epigenetic clocks and prevalent disease as well as an assessment of their ability to predict time-to-disease onset. These findings may expedite the future use and refinement of large-scale molecular data-based approaches for predicting clinically-defined outcomes and subsequent individual disease risk prediction.

## Results

### Demographics and Epigenetic Clocks

In the discovery cohort, 56.3% of the participants were female with a mean age of 51.4 years (SD: 13.2) (n = 4,450). The mean values for epigenetic measures were as follows: HorvathAge (60.1 years, SD: 9.8), HannumAge (47.4 years, SD: 9.6), DNAm PhenoAge (43.7 years, SD: 11.5), DNAm GrimAge (48.8 years, SD: 10.9) and DNAm Telomere Length (7.4 kilobase pairs, SD: 0.3). Summary data for all variables in this study are presented in Supplementary File 1.

In the replication cohort, 61.4% were female with a mean age of 50.0 years (SD: 12.5) (n = 2,578). Values for all phenotypes were comparable between discovery and replication cohorts with the exception of DNAm GrimAge (discovery: 48.8 years, SD: 10.9, replication: 60.5 years, SD: 10.6), and the incidence of self-reported depression (discovery: 8.34%, replication: 16.36%), and SCID (Structured Clinical Interview for DSM)-identified Depression (discovery: 18.54%, replication: 38.17%). This reflects ascertainment bias in the replication cohort in which there was enrichment for depression and may have contributed to the higher DNAm GrimAge when compared to the discovery cohort.

### Epigenetic Clocks and Disease Prevalence

In a basic model adjusting for age and sex, 37 phenotypes were significant at Bonferroni-corrected levels of significance in both the discovery and replication cohorts (Supplementary Note 1; Supplementary Tables 1-4). Supplementary Figures 1, 2 and 3 highlight significant associations for categorical traits, continuous traits and all-cause mortality, respectively. A clock-by-clock comparison of associations with categorical and continuous phenotypes from fully-adjusted models in the replication cohort, stratified by disease type, is shown in Supplementary File 2. For all models, beta coefficients for continuous traits were correlated 0.95 between discovery and replication sets. For categorical phenotypes, the correlation coefficient for log odds was 0.83 between sets (Supplementary Figure 5).

Twelve relationships remained significant in both discovery and replication sets in a fully-adjusted model accounting for age, sex and five common risk factors (Supplementary Tables 5 and 6, respectively). Those relationships which were significant in both cohorts at P < 8.20 × 10^-4^ are reported herein and presented in Table 1 and Figure 1.

**Figure 1.**
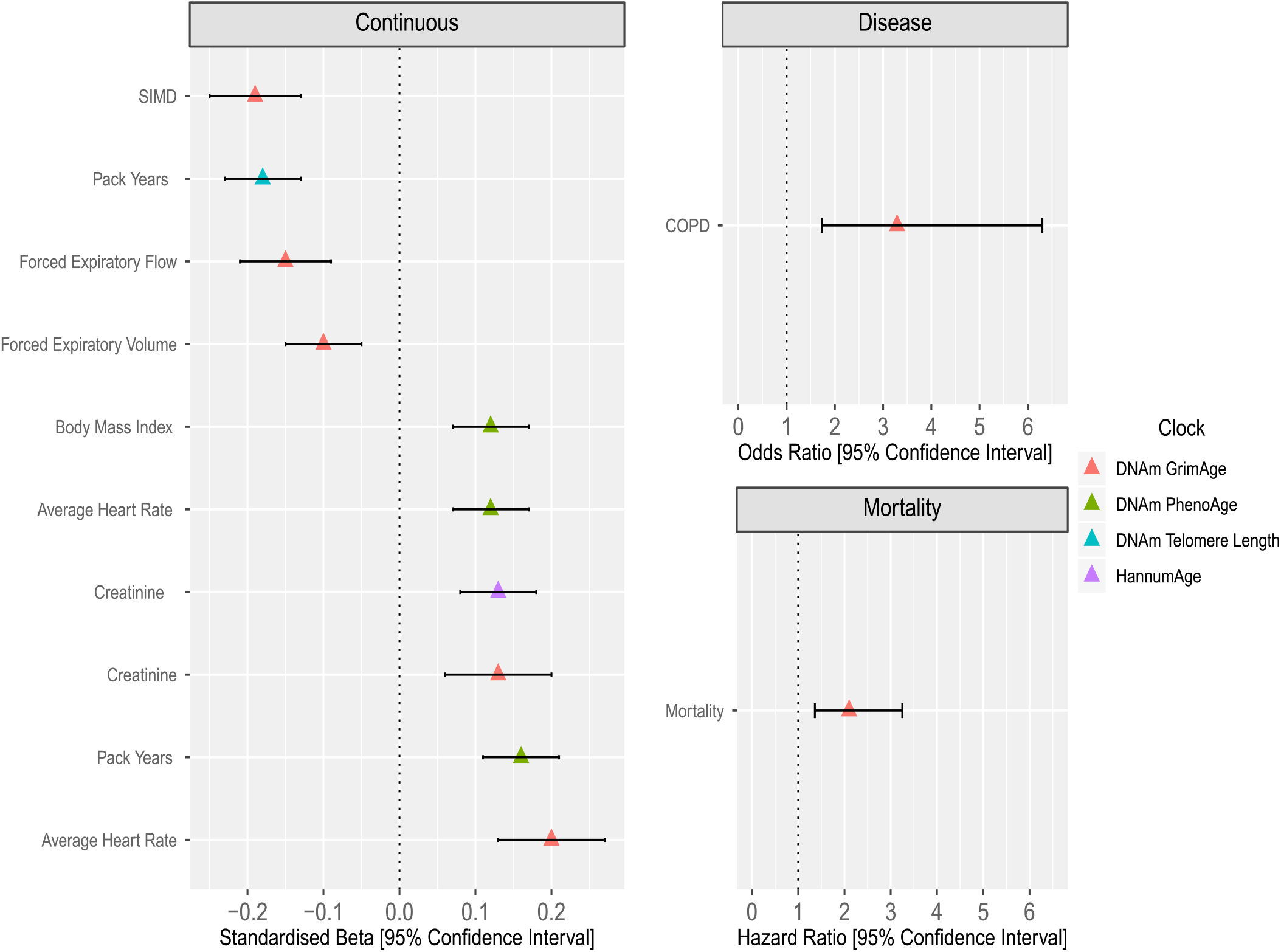
The association of epigenetic clocks with prevalence of common disease, associated continuous traits and all-cause mortality in Generation Scotland in a model adjusting for age, sex and common disease risk factors. Only associations present in discovery and replication sets are shown, and replication test statistics are presented. *Continuous:* Age-adjusted DNAm GrimAge was associated with greater deprivation (lower SIMD), reduced forced expiratory flow and forced expiratory volume. Age-adjusted DNAm GrimAge was positively associated with creatinine levels and average heart rate. Age-adjusted DNAm PhenoAge was positively associated with body mass index, average heart rate and smoking pack years. Age-adjusted DNAm Telomere Length was negatively associated with smoking pack years. Age-adjusted HannumAge was positively associated with creatinine levels. *Disease:* Age-adjusted DNAm GrimAge alone was associated with prevalence of COPD in both discovery and replication sub-cohorts after correction for multiple testing. *All-Cause Mortality:* Age-adjusted DNAm GrimAge alone was associated with all-cause mortality in both test sets after multiple testing correction. Associations represent a one standard deviation increase in the respective age-adjusted epigenetic clock measures. Models were adjusted for age, sex, alcohol, body mass index, deprivation, education and smoking. Models involving lung function tests were also corrected for height. COPD (chronic obstructive pulmonary disease), SIMD (Scottish Index of Multiple Deprivation).

**Table 1.**
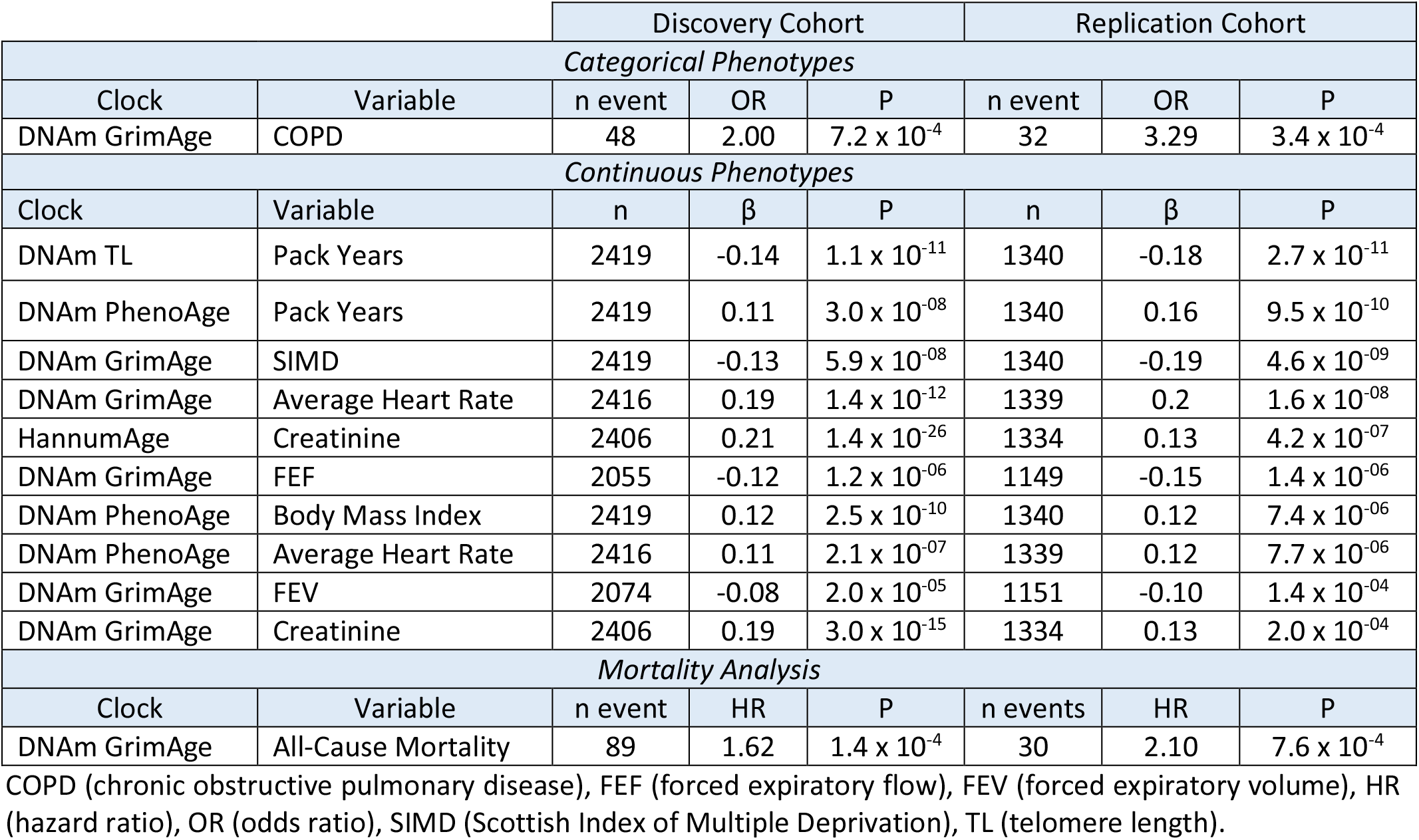
Significant relationships between epigenetic clocks and prevalent disease data, with related continuous traits, present in both discovery and replication cohorts. Analyses were performed using a fully-adjusted model accounting for age, sex, alcohol consumption, body mass index, deprivation, education and smoking pack years.

### Associations with disease

In relation to self-report disease data, only the association between an accelerated DNAm GrimAge and COPD remained significant in both cohorts in the fully-adjusted model (replication cohort: OR per SD = 3.29, 95% Confidence Interval (CI) = [1.73, 6.30], P = 3.4 × 10^-4^; Figure 1).

### Associations with All-Cause Mortality

An accelerated DNAm GrimAge alone was associated with all-cause mortality following adjustment for the lifestyle risk factors (replication cohort: HR per SD = 2.10, 95% CI = [1.36, 3.25], P = 7.6 × 10^-4^; Figure 1).

### Associations with Continuous Clinically-Associated Traits

An accelerated DNAm GrimAge was associated with greater deprivation (SIMD; β_replication_ = −0.19, 95% CI = [-0.25, −0.13], P = 4.6 × 10^-9^), an increased average heart rate (β_replication_ = 0.20, 95% CI = [0.13, 0.27], P = 1.6 × 10^-8^), a reduced forced expiratory flow (β_replication_ = −0.15, 95% CI = [-0.21, −0.09], P = 1.4 × 10^-6^), a reduced forced expiratory volume (β_replication_ = −0.10, 95% CI = [-0.15, −0.05], P = 1.4 × 10^-4^) and increased creatinine levels (β_replication_ = 0.13, 95% CI = [0.06, 0.20], P = 2.0 × 10^-4^).

An accelerated DNAm PhenoAge was positively associated with smoking pack years (replication cohort: β_replication_ = 0.16, 95% CI = [0.11, 0.21], P = 9.5 × 10^-10^), an increased body mass index (β_replication_ = 0.12, 95% CI = [0.07, 0.17], P = 7.4 × 10^-6^) and an increased average heart rate (β_replication_ = 0.12, 95% CI = [0.07, 0.17], P = 7.7 × 10^-6^).

Age adjusted DNAm Telomere Length was negatively associated with smoking pack years (β_replication_ = −0.18, 95% CI = [-0.23, −0.13], P = 2.7 × 10^-11^). An accelerated DNAm HannumAge (EEAA) was associated with increased creatinine (β_replication_ = 0.13, 95% CI = [0.08, 0.18], P = 4.2 × 10^-7^).

### Covariate-specific attenuation

To examine the contribution of the five risk factors to attenuating the 37 significant associations tested above, we repeated each model including only one of these five covariates at a time. On average, the five factors displayed similar degrees of mean attenuation of tested traits (discovery: range = [9.2%, 12.6%], replication: range = [5.7%, 16.4%]; Supplementary Tables 7 and 8, respectively). Smoking pack years exhibited the highest mean attenuation of trait-epigenetic clock relationships in both cohorts (discovery: 12.6%, replication: 16.4%).

### Epigenetic Clocks and Disease Incidence

For incident disease outcomes, there were 13 Bonferroni-corrected significant associations (Supplementary Note 2, Supplementary Figure 4, full output in Supplementary Table 9). Of these, 5 remained significant in a fully-adjusted model at a Bonferroni-corrected significance threshold of 1 × 10^-3^ (Supplementary Table 10). These relationships are presented herein and in Figure 2.

**Figure 2.**
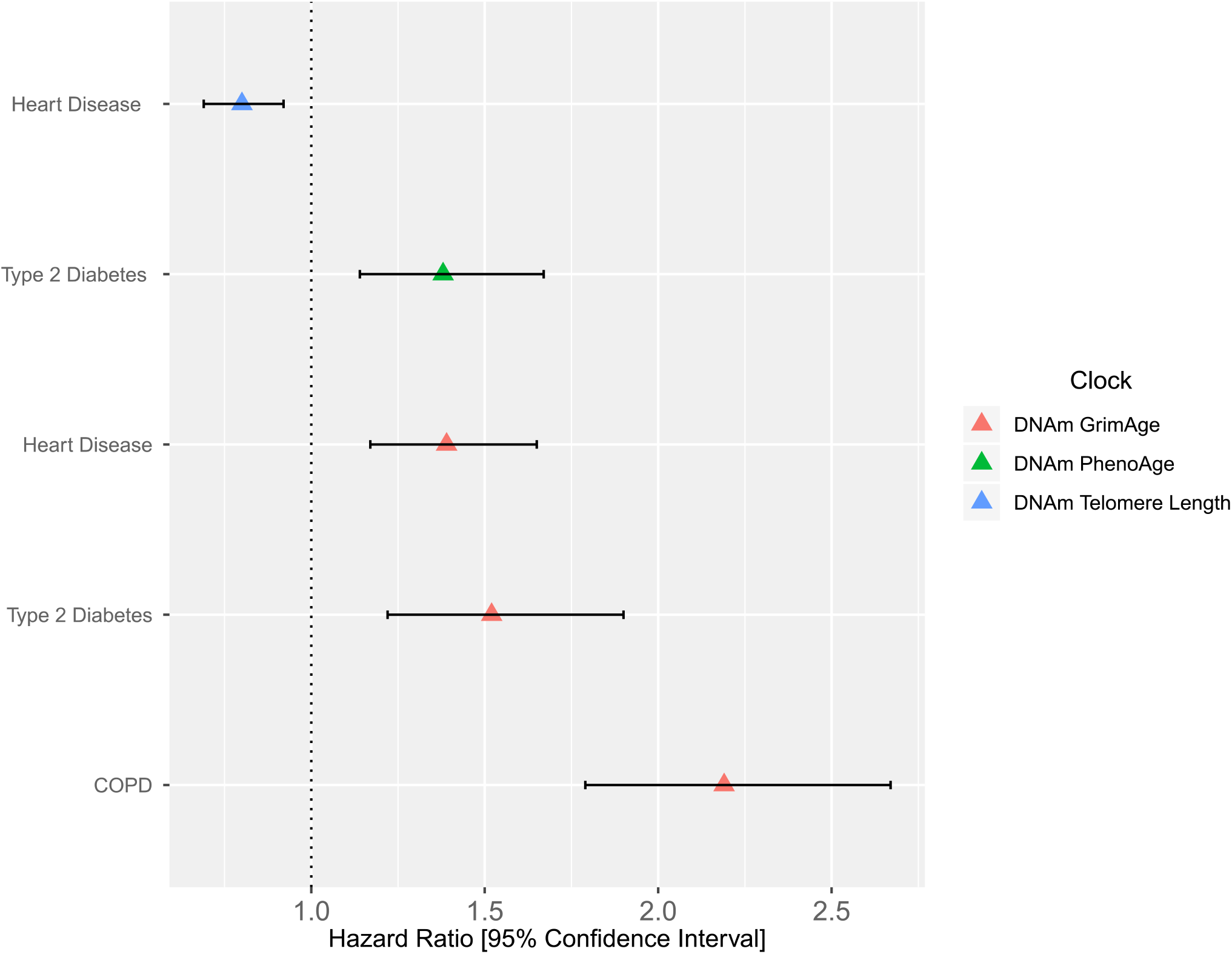
The association of epigenetic clocks with incidence of disease in Generation Scotland in a model adjusting for age, sex and common disease risk factors. Age-adjusted DNAm GrimAge was associated with incidence of COPD, type 2 diabetes and ischemic heart disease after thirteen years of follow-up. Age-adjusted DNAm PhenoAge associated with incidence of type 2 diabetes. Age-adjusted measures of DNAm Telomere Length associated with incidence of ischemic heart disease. Associations represent a one standard deviation increase in age-adjusted epigenetic clocks. Models were adjusted for age, sex, alcohol, body mass index, deprivation, education and smoking. COPD (chronic obstructive pulmonary disease).

A one standard deviation increase in DNAm GrimAge at baseline was associated with incidence of COPD (HR = 2.19, 95% CI = [1.79, 2.67], P = 1.9 × 10^-14^), type 2 diabetes (HR = 1.52, 95% CI = [1.22, 1.90], P = 8.9 × 10^-12^) and heart disease (HR = 1.39, 95% CI = [1.17, 1.65], P = 1.4 × 10^-4^). An accelerated DNAm PhenoAge (per SD) associated with a higher incidence of type 2 diabetes (HR = 1.39, 95% CI = [1.14, 1.67], P = 9.9 × 10^-4^). Age-adjusted DNAm Telomere Length (per SD) associated with a lower incidence of heart disease (HR = 0.80, 95% CI = [0.69, 0.92], P = 2.5 × 10^-4^).

### Sex-specific analysis of epigenetic clocks and phenotypes in Generation Scotland

As the occurrence of common diseases differs between the sexes, we ran a sensitivity analysis using prevalence data to determine the correlation between effect sizes of trait-clock comparisons for males versus females. In the discovery cohort, continuous phenotypes possessed a correlation coefficient of 0.92 between sexes whereas categorical disease phenotypes exhibited a correlation coefficient of 0.81 (Supplementary Figure 5). In the replication cohort, there was a correlation of 0.84 and 0.65 between effect sizes for continuous and categorical phenotypes, respectively (Supplementary Figure 7). Excluding diseases ≤ 10 cases (lung and bowel cancer), the largest difference between males and females was for the IEAA-stroke relationship (males: no. of events = 35, OR = 1.15; females: no. of events = 24, OR = 0.79). On average, the largest difference between males and females across clocks was observed for COPD with males having a higher odds ratio for each clock (mean difference in effect sizes across clocks = 0.18, range = [0.12, 0.37]) (discovery cohort; Supplementary Table 11).

## Discussion

In this study, we examined the associations between five major epigenetic clocks and the prevalence and incidence of the leading causes of mortality and disease burden in high-income countries. DNAm GrimAge, a predictor of mortality, associated with the prevalence of COPD and incidence of various disease states, including COPD, type 2 diabetes and cardiovascular disease. It was associated with death due to all-cause mortality and outperformed competitor clocks in capturing variability in clinically-associated continuous traits. A higher-than-expected DNAm PhenoAge predicted incidence of type 2 diabetes in the present study. Age-adjusted measures of DNAm Telomere Length associated with incidence of ischemic heart disease. Our results replicate previous cross-sectional findings between DNAm PhenoAge and BMI, diabetes (19) and socioeconomic position (in a basic model) (26). We also replicated associations between DNAm GrimAge and heart disease (11). Lastly, we replicated the relationship of HannumAge with creatinine (28) and of DNAmTLadjAge with smoking pack years (29).

DNAm GrimAge served as a powerful correlate of various phenotypes in our study, and has been previously shown to associate with incident heart disease, time-to-cancer and neurological health (14, 20). DNAm GrimAge is derived from chronological age, sex and methylation-based surrogates of smoking pack years and seven plasma proteins (including DNAm-based estimators of plasminogen activator inhibitor 1, growth differentiation factor 15 and cystatin C). Here, we show that this blood-based epigenetic predictor of mortality risk is associated with poorer performance in lung function tests and predicted incidence of COPD. Compromised lung function previously has been linked to mortality (30, 31). While it is possible that the associations are mainly driven by the inclusion of smoking pack years, DNAm GrimAge remained associated with COPD and spirometry tests when controlling for self-reported smoking pack years. DNAm PhenoAge predicted incidence of type 2 diabetes; however, this may reflect the inclusion of HbA1c in the phenotypic age measure which is used to diagnose diabetes. In our study, an epigenetic predictor of telomere length predicted time-to-onset of ischemic heart disease. A shorter leukocyte telomere length has been shown to associate with heart disease in diverse populations, suggesting that the DNAm Telomere Length predictor may capture key facets of this clinical association (32–34). Our rich resource of genome-wide DNA methylation and longitudinal health data is the first to show the association of epigenetic clocks with a wide range of common disease states, even after accounting for major confounding influences. These findings have implications for the potential utility of epigenetic clocks in clinical settings.

The majority of our prevalent disease data relied on self-report. Self-report prevalence data has been shown to have a high degree of sensitivity and specificity (35). Our incident data was obtained using ICD-10 codes from health record linkage. Strikingly, epigenetic clocks showed strong associations with the incidence of common diseases following thirteen years of follow-up from study baseline. These clocks performed better at predicting incident rather than prevalent data. However, this may reflect the inclusion of health record-linked versus self-report data, and the larger sample size in incidence analyses.

An important limitation is the lack of adjustments for medication use, which may confound associations between clocks and chronic conditions. Furthermore, studies examining causality between the relationships shown are merited. It is also unclear whether the risk factors examined in this study play a casual role in driving associations between epigenetic clocks and phenotypes, or whether these pleiotropically affect both altered DNA methylation and adverse health outcomes. Genetic influences may contribute to differences in DNA methylation and the subsequent estimation of epigenetic age; therefore, it is possible that our findings may not be generalisable to individuals of non-European ancestry (36, 37).

In summary, using a large cohort with rich health and epigenetic data, we provide the first across-five-epigenetic-clocks comparison of epigenetic predictors with respect to leading causes of mortality and disease burden. DNAm GrimAge outperformed the other clocks in its associations with disease data and associated clinical traits. This may suggest that predicting mortality, rather than age or homeostatic characteristics, may be more informative for common disease prediction. Thus, proteomic-based methods (as utilised by DNAm GrimAge) using large, physiologically diverse protein sets for predicting ageing and health may be of particular interest in future studies. Our results may help to refine the future use and development of ageing biomarkers, particularly in studies which aim to comprehensively examine their ability to predict stringent clinically-defined outcomes. Our analyses suggest that epigenetic clocks can predict incidence of common disease states, even after accounting for major confounding risk factors. This may have significant implications for their potential utility in clinical settings to complement gold-standard methods of clinical disease assessment and management.

## Materials and Methods

### Generation Scotland

Details of the Generation Scotland (GS) study have been described previously (38, 39). Briefly, the cohort includes 23,960 individuals, most (94.16%) with at least one other first-degree family member participating in the study. This encompasses 5,573 families with a median family size of 3 (interquartile range: 2 – 5 members; excluding 1,400 singletons without any relatives in the study). For prevalence analyses, the discovery cohort comprised unrelated GS participants with genome-wide methylation data (n_discovery_ = 4,450). The replication cohort was also derived from GS participants, unrelated to those in the discovery cohort, who had genome-wide DNA methylation measured in a separate batch (n_replication_ = 2,578). Within the replication cohort, participants were also unrelated to one another. For incidence analyses, all individuals with available methylation and phenotypic data in GS were considered (n = 9,537).

### DNA Methylation and Estimation of Clocks

DNA methylation levels were measured using the Illumina HumanMethylationEPIC BeadChip Array on blood samples from GS participants. Further details on the processing of methylation data and the calculation of the five clocks are outlined in Supplementary Methods; all five clocks were calculated using Horvath’s online age calculator (https://dnamage.genetics.ucla.edu/). Normalised GS methylation data were uploaded as input for the algorithm. Data underwent a further round of normalisation by the age calculator. Briefly, HorvathAge provides an estimate of biological ageing termed “intrinsic epigenetic age acceleration (IEAA)” as it is independent of age-related changes in blood composition. IEAA is derived from regressing HorvathAge onto chronological age. In contrast, HannumAge provides a measure of ageing referred to as “extrinsic epigenetic age acceleration (EEAA)” as it encompasses age-related changes in blood cell composition. EEAA is derived from regressing a weighted average of HannumAge and three blood cell types (naive and exhausted cytotoxic T-cells, and plasmablasts) onto chronological age. DNAm PhenoAge reflects an individual’s ‘Phenotypic Age’ and, when regressed onto chronological age, provides an index of age acceleration termed ‘AgeAccelPheno’. Similarly, when age-adjusted, DNAm GrimAge is termed ‘AgeAccelGrim’. Lastly, age-adjusted ‘DNAm Telomere Length’ is referred to as ‘DNAmTLadjAge’. These fives measures of biological age acceleration were input as independent variables in statistical models. Correlations between these predictors are shown in Supplementary Figure 8. DNAmTLadjAge was negatively correlated with the other four indices of age acceleration (mean coefficient = −0.36, range = −0.12 to −0.47). This negative correlation is present as shorter telomere lengths typically correspond to an advanced age. The mean correlation coefficient between the remaining four predictors was 0.34 (range = 0.11 to 0.50).

### Phenotype Preparation

For continuous phenotypes, outliers were defined as those values which were beyond 3.5 standard deviations from the mean for a given trait. These outliers were removed prior to analyses. Body mass index was log transformed. To reduce skewness in the distribution of alcohol consumption and smoking pack years, a log(units + 1) or log(pack years + 1) transformation was performed. The interval from the start of the Q wave to the end of the T wave on electrocardiogram tests (QT interval) was corrected for heart rate. A general fluid (‘*gf*’) cognitive ability score was derived from principal components analysis of three tests examining different cognitive domains. These domains were processing speed (Wechsler Digit Symbol Substitution Test), verbal declarative memory (Wechsler Logical Memory Test) and verbal fluency (the phonemic Verbal Fluency Test). To derive a general (‘*g’*) cognitive ability score, principal component analysis was performed on the above three tests and a measure of crystallised intelligence: The Mill Hill Vocabulary test. The first unrotated principal components from these analyses were extracted and labelled as ‘*gf*’ and ‘*g*’, respectively.

For categorical phenotypes, we aimed to examine the ten leading causes of mortality in high-income countries (40). We also aimed to examine the ten leading causes of disease burden, six of which overlap with the top causes of mortality. This represents fourteen diseases. We had self-report phenotypic information for the prevalence of nine of these diseases (Supplementary File 1); specifically, we lacked self-report information on lower respiratory diseases and kidney disease (mortality), skin and sense organ diseases (disease burden), and Alzheimer’s disease (AD). We were able to use proxy phenotypes for two of these conditions. We used self-reported maternal history and paternal history as proxies for AD. For kidney disease, we estimated glomerular filtration rate (eGFR) from serum creatinine levels using the Chronic Kidney Disease Epidemiology Collaboration CKD-EPI equation (41) from which we inferred the prevalence of chronic kidney disease (CKD). Individuals with an eGFR < 60 ml/min/1.73 m^2^ were considered to have CKD. In addition to self-report depression, we also had available information on SCID (Structured Clinical Interview for DSM)-identified depression (42). Lastly, we separated self-reported back and neck pain into distinct phenotypes for analyses. Together, this resulted in a total of fourteen disease phenotypes for prevalence analyses.

In relation to disease incidence, health record linkage was available for up to thirteen years of follow-up since study baseline (median time-of-onset from baseline = 5.75 years, range = [<1 month, 13 years]). For each disease state, those individuals who self-reported disease at study baseline were excluded. For cancer, individuals present on the Scottish Cancer Registry (SMR06) were included as cases for incidence analyses. Additionally, for incident cancer analyses, individuals who were recorded on the General Acute Inpatient and Day Case - Scottish Morbidity Record (SMR01) were removed from the control set. For a given condition, individuals who self-reported no disease at study baseline but had prior evidence of diagnosis through health record linkage were removed from analyses. Discovery and replication cohorts were combined to consider all participants for follow-up and to provide sufficient number of cases for analyses. For incident disease analyses, ICD-10-coded data were retrieved for the following ten conditions: AD, bowel cancer, breast cancer, COPD, depression, type 2 diabetes, dorsalgia (neck and back pain combined), ischemic heart disease, lung cancer and stroke. These reflect the disease states examined in the prevalence analyses with the exception of kidney disease. Furthermore, the two proxies of AD, two measures of depression and separate measures of neck and back pain were replaced by single, clinically-defined counterparts in the incidence analyses.

### Statistical Analysis

Age acceleration was defined as the residual term from regressing an epigenetic predictor onto chronological age. Linear regression models were used to examine the association between continuous traits and age acceleration. In cross-sectional analyses, logistic regression was used to test the association between categorical disease phenotypes and age acceleration. In longitudinal analyses, Cox proportional hazards regression models were used to examine whether measures of biological age were associated with incidence of disease. Cox models were also used to examine whether age-adjusted epigenetic clocks were associated with all-cause mortality in discovery and replication cohorts. The proportional hazards assumption was tested using the *cox.zph()* function in the *survival* package in *R.* There was no strong evidence (P > 0.05) of assumption violation for the reported significant associations. Phenotypes were scaled to mean zero and unit variance. Continuous or categorical phenotypes were input as dependent variables with biological clocks incorporated as independent variables.

In a basic model, all analyses were adjusted for chronological age and sex. Additional adjustments for height and smoking status were carried out for measures of lung function. All significant tests from the basic model were then repeated adjusting for an additional five covariates, which represent important risk factors for common diseases. These covariates were: alcohol intake (units consumed/week), body mass index, educational attainment, deprivation (Scottish Index of Multiple Deprivation) and tobacco smoking pack years.

> ***Basic model**: Phenotype ~ Biological Clock + age + sex*
>
> ***Fully-adjusted model**: Phenotype ~ Biological Clock + age + sex + alcohol units consumed per week + body mass index + educational attainment + Scottish Index of Multiple Deprivation + smoking pack years*

In relation to cross-sectional prevalence data, the discovery analyses consisted of 33 phenotypes which were tested against every clock (all-cause mortality, fourteen disease and eighteen continuous phenotypes; Supplementary Table 1). This led to a total of 165 (33 × 5 clock) tests; however, the DNAm GrimAge versus smoking pack years comparison was excluded given the inclusion of a methylation-based surrogate of pack years in the development of DNAm GrimAge. This led to a Bonferroni-corrected significance threshold of P<0.05/164 tests = 3.05 × 10^-4^. Of these 164 tests, 61 were significant; thus, in the replication cohort, a Bonferroni-corrected significance threshold of P<0.05/61 tests = 8.20 × 10^-4^ was set. In total, 37 associations were significant in both cohorts. The fully-adjusted model was then applied to these associations in both the discovery and replication cohorts, with the same Bonferroni-corrected threshold of P < 0.05/61 = 8.20 × 10^-4^.

In relation to incidence data, all ten phenotypes were tested against each of the five clocks. In the initial basic model, this resulted in a Bonferroni-corrected significance threshold of P<0.05/50 tests = 1.0 × 10^-3^. In total, thirteen associations were significant and brought forward to the fully-adjusted analysis stage. In the fully-adjusted model, the same Bonferroni-corrected significance threshold of P<0.05/50 tests = 1.0 × 10^-3^ was applied.

### Ethics

GS has been ethically approved and granted research tissue bank status by the NHS Tayside Committee on Medical Research Ethics (REC reference numbers: 05/S1401/89 and 10/S1402/20, respectively).

## Acknowledgements

GS received core support from the Chief Scientist Office of the Scottish Government Health Directorates (CZD/16/6) and the Scottish Funding Council (HR03006). DNA methylation profiling of the GS samples was carried out by the Genetics Core Laboratory at the Wellcome Trust Clinical Research Facility, Edinburgh, Scotland and was funded by the Medical Research Council UK, the Brain & Behavior Research Foundation (Ref: 27404) and the Wellcome Trust (Wellcome Trust Strategic Award “STratifying Resilience and Depression Longitudinally” ((STRADL) Reference 104036/Z/14/Z)). AMM is supported by the Wellcome Trust (104036/Z/14/Z, 216767/Z/19/Z), UKRI MRC (MC_PC_17209, MR/S035818/1) and the European Union H2020 (SEP-210574971). AJS and RFH are supported by funding from the Wellcome Trust 4-year PhD in Translational Neuroscience – training the next generation of basic neuroscientists to embrace clinical research [RFH: 108890/Z/15/Z; AJS: 203771/Z/16/Z]. DLMcC and REM are supported by Alzheimer’s Research UK major project grant ARUK-PG2017B-10. SH is supported by 1U01AG060908 –01. DMH is supported by a Sir Henry Wellcome Postdoctoral Fellowship (Reference 213674/Z/18/Z) and a 2018 NARSAD Young Investigator Grant from the Brain & Behavior Research Foundation (Ref: 27404).

