## Supplementary Figures for "Epigenetic clocks predict prevalence and incidence of leading causes of death and disease burden"

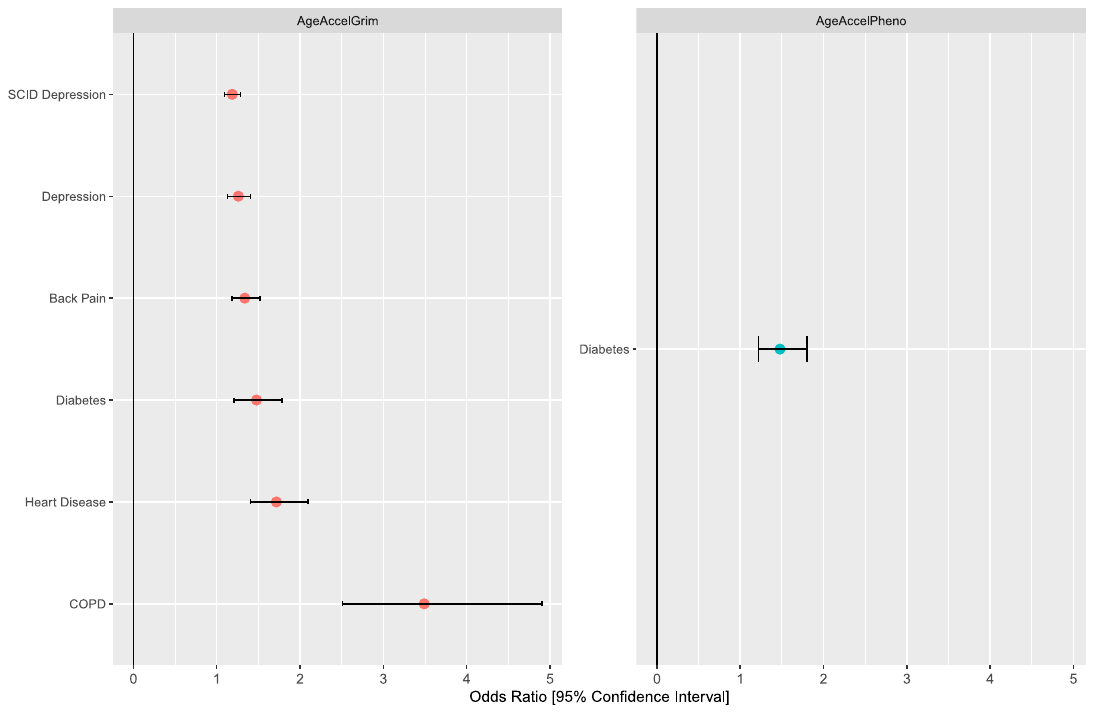

**Supplementary Figure 1. The association of epigenetic clocks with disease phenotypes in Generation Scotland.** A higher-than-expected DNAm GrimAge (higher AgeAccelGrim) was positively associated with SCID depression and self-reported measures of depression, back pain, diabetes, heart disease and COPD. A higher-than-expected DNAm PhenoAge (higher AgeAccelPheno) was positively associated with diabetes. COPD (chronic obstructive pulmonary disease), SCID (Structured Clinical Interview for DSM).

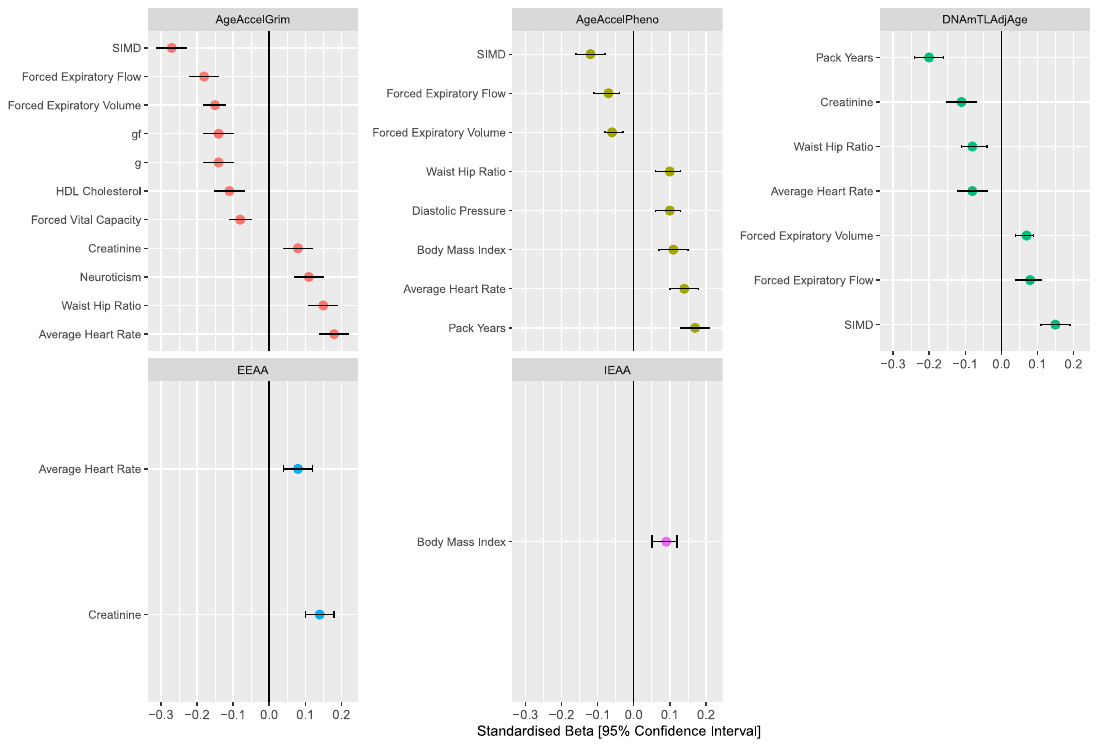

**Supplementary Figure 2. The association of epigenetic clocks with continuous traits in Generation Scotland.** AgeAccelGrim (DNAm GrimAge) was negatively associated with SIMD, forced expiratory flow, forced expiratory volume, a general factor of fluid intelligence (gf), a general factor of cognitive ability (g), HDL cholesterol and forced vital capacity. AgeAccelGrim was positively associated with creatinine, neuroticism, waist-to-hip ratio and average heart rate. AgeAccelPheno (DNAm PhenoAge) was negatively associated with SIMD, forced expiratory flow and forced expiratory volume. AgeAccelPheno was positively associated with waist-to-hip ratio, diastolic pressure, body mass index, average heart rate and smoking pack years. DNAmTLadjAge (DNAm Telomere Length) was negatively associated with smoking pack years, creatinine, waist-to-hip ratio and average heart rate. DNAmTLadjAge was positively associated with forced expiratory volume, forced expiratory flow and SIMD. EEAA (HannumAge) was positively associated with average heart rate and creatinine. IEAA (HorvathAge) was positively associated with body mass index. HDL (high-density lipoprotein), SIMD (Scottish Index of Multiple Deprivation).

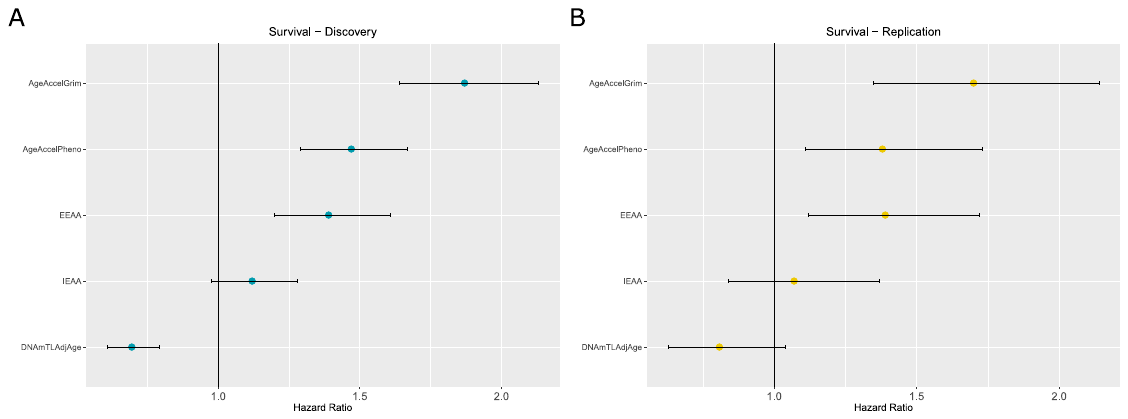

**Supplementary Figure 3. The association of epigenetic clocks with all-cause mortality in Generation Scotland.** Estimates for biological age were regressed onto chronological age to provide a measure of biological age acceleration. In the discovery and replication cohorts, there were 182 (4.09%) and 57 (2.21%) deaths, respectively. Following correction for multiple testing, AgeAccelGrim (DNAm GrimAge) alone was significantly associated with all-cause mortality in the discovery (n = 4,450) and replication (n = 2,578) cohorts.

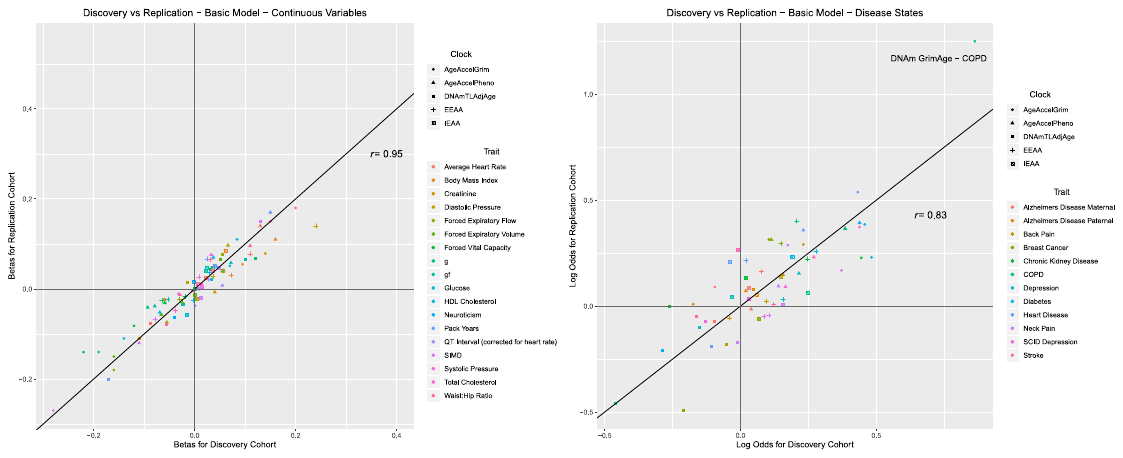

**Supplementary Figure 4. Degree of correlation between continuous variables (A), or categorical variables (B), across discovery and replication cohorts.** Lung and bowel cancer were excluded due to a low number of cases < 10.

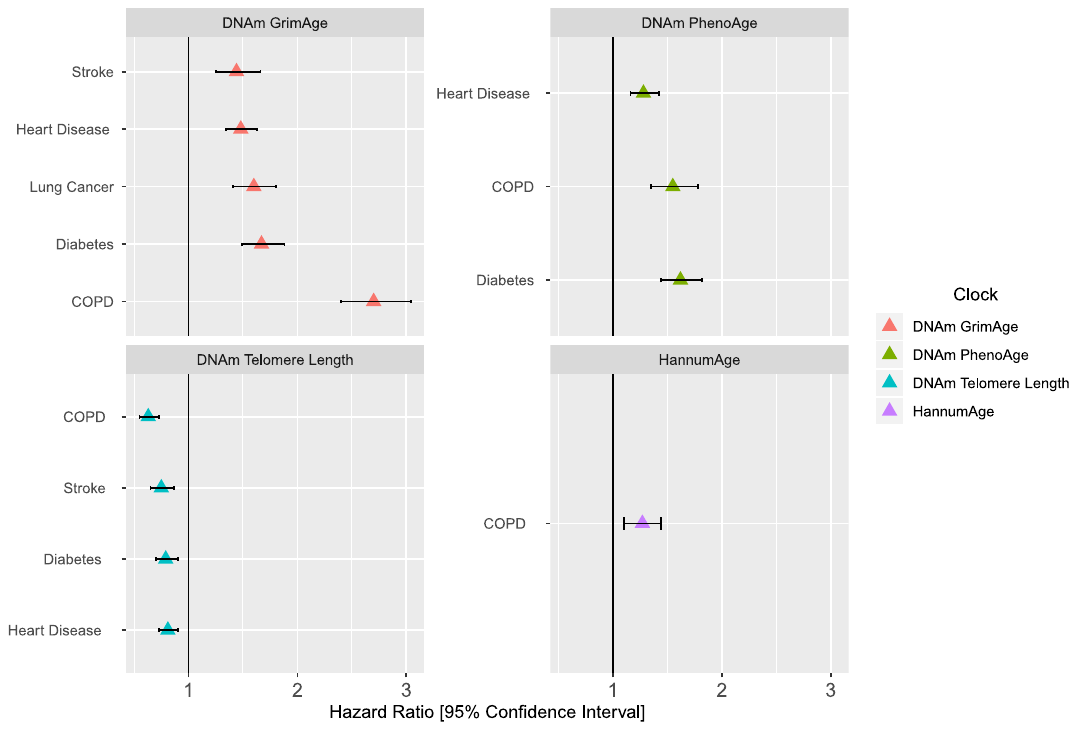

**Supplementary Figure 5. The association of epigenetic clocks with incidence of disease in Generation Scotland in a basic model adjusting only for age and sex.** Age-adjusted DNAm GrimAge was associated with incidence of COPD, diabetes, lung cancer, ischemic heart disease and stroke after thirteen years of follow-up. Age-adjusted DNAm PhenoAge associated with incidence of diabetes, COPD and ischemic heart disease. Age-adjusted measures of DNAm Telomere Length associated with incidence of heart disease, diabetes, stroke and COPD. Age-adjusted HannumAge associated only with incident COPD. COPD (chronic obstructive pulmonary disease).

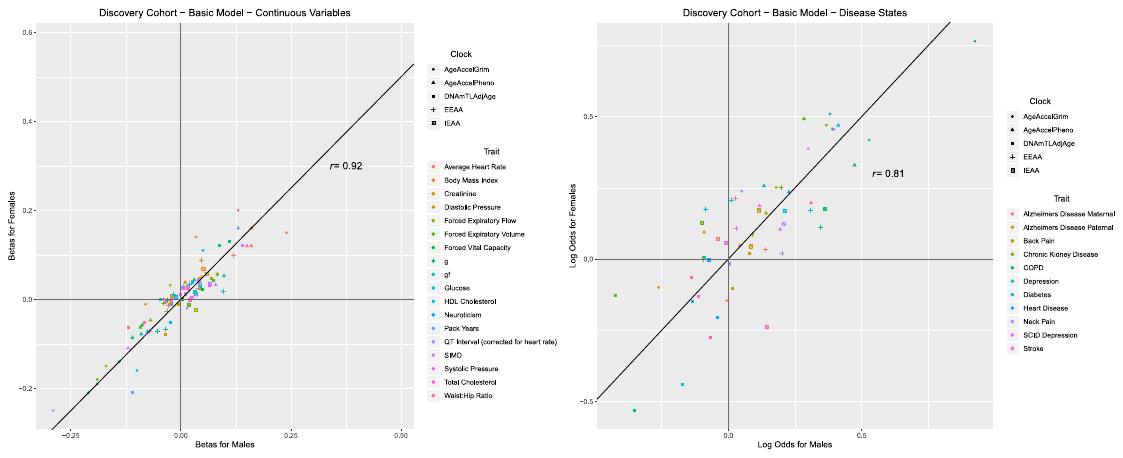

**Supplementary Figure 6. Degree of correlation for continuous variables (A), or categorical variables (B), between males and females in discovery cohort.** Lung and bowel cancer were excluded due to a low number of cases < 10. Breast cancer was excluded as only female cases were present in Generation Scotland.

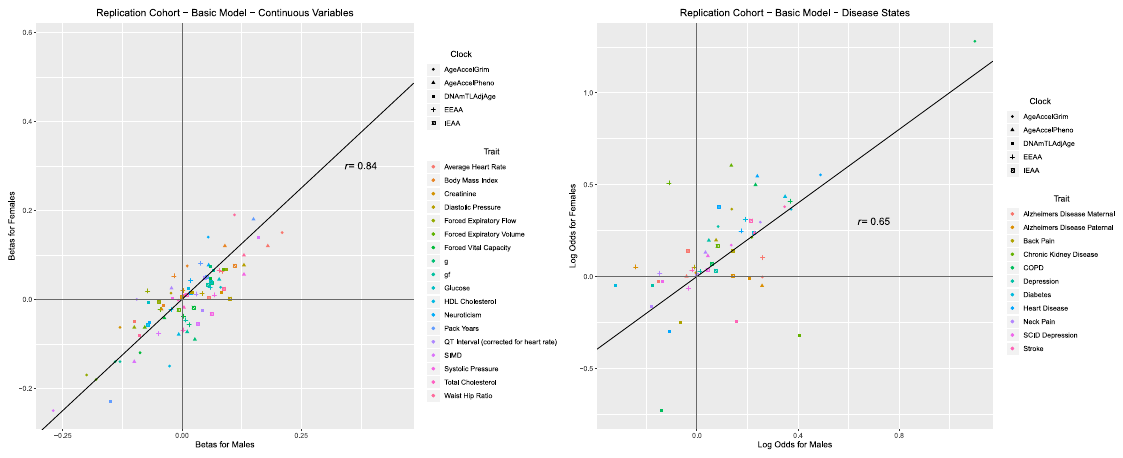

**Supplementary Figure 7. Degree of correlation for continuous variables (A), or categorical variables (B), between males and females in replication cohort.** Lung and bowel cancer were excluded due to a low number of cases < 10. Breast cancer was excluded as only female cases were present in Generation Scotland.

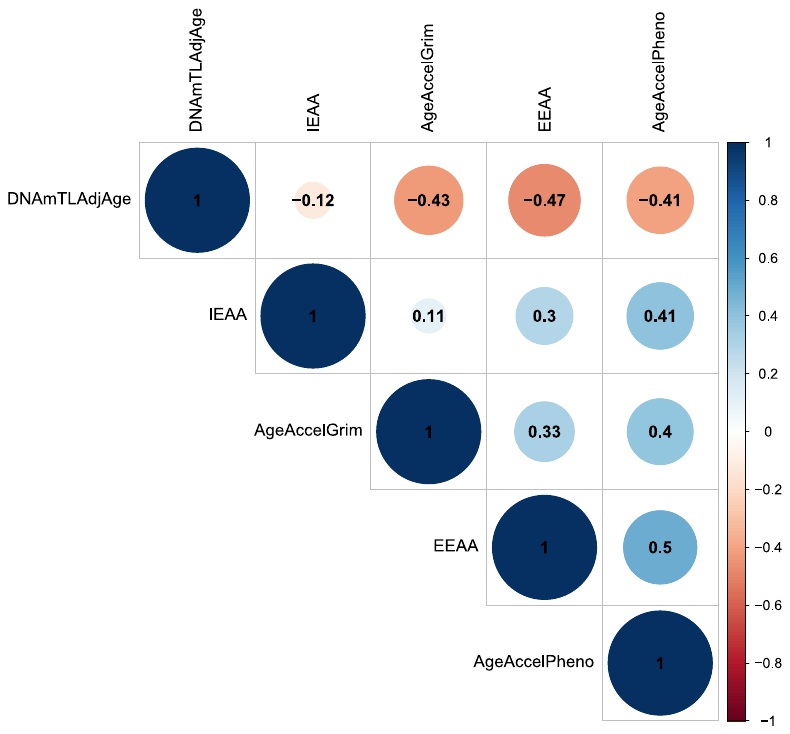

**Supplementary Figure 8. Correlation among different epigenetic measures of biological age.** All age-adjusted measures of biological clocks are positively correlated, with the exception of age-adjusted DNAm Telomere Length (DNAmTLAdjAge). AgeAccelGrim and AgeAccelPheno refer to age-adjusted DNAm GrimAge and DNAm PhenoAge, respectively. EEAA (HannumAge), IEAA (HorvathAge).
