## Supplementary File 1 for "Epigenetic clocks predict prevalence and incidence of leading causes of death and disease burden"

Supplementary File 1. Demographics and Descriptive Statistics for Discovery and Replication Cohorts.

| | ***Discovery*** | | | ***Replication*** | | |
|---|---|---|---|---|---|---|
| *Variable* | *n* | *Mean* | *SD* | *n* | *Mean* | *SD* |
| Age (years) | 4450 | 51.40 | 13.20 | 2578 | 50.00 | 12.50 |
| | *n* | *Females/Males* | *%Female* | *n* | *Females/Males* | *%Females* |
| Sex | 4450 | 2506/1944 | 56.31% | 2578 | 1583/995 | 61.40% |
| *Epigenetic Clocks* | *n* | *Mean* | *SD* | *n* | *Mean* | *SD* |
| DNAm GrimAge (years) | 4450 | 48.82 | 10.89 | 2578 | 60.47 | 10.63 |
| DNAm PhenoAge (years) | 4450 | 43.73 | 11.45 | 2578 | 44.98 | 9.75 |
| Horvath Age (EEAA; years) | 4450 | 60.13 | 9.83 | 2578 | 54.75 | 9.35 |
| Hannum Age (IEAA; years) | 4450 | 47.38 | 9.64 | 2578 | 46.97 | 9.13 |
| DNAm Telomere Length (TL; kbp) | 4450 | 7.42 | 0.26 | 2578 | 7.31 | 0.26 |
| *Continuous Phenotypes* | *n* | *Mean* | *SD* | *n* | *Mean* | *SD* |
| Average Heart Rate (beats/min) | 4444 | 69.20 | 11.40 | 2572 | 69.70 | 11.10 |
| Body Mass Index (kg/m$^2$) | 4423 | 26.80 | 4.96 | 2567 | 27.20 | 5.35 |
| Creatinine (µmol/L) | 4427 | 71.20 | 14.60 | 2566 | 70.10 | 14.50 |
| Diastolic Pressure (mmHg) | 4447 | 80.80 | 10.50 | 2573 | 80.10 | 9.93 |
| Forced Expiratory Flow (L/s) | 3750 | 2.99 | 1.23 | 2191 | 2.99 | 1.21 |
| Forced Expiratory Volume (L) | 3776 | 3.02 | 0.84 | 2191 | 3.00 | 0.82 |
| Forced Vital Capacity (L) | 3775 | 3.97 | 0.99 | 2197 | 3.93 | 0.97 |
| General Factor of Fluid Intelligence | 4324 | -0.06 | 0.99 | 2529 | 0.02 | 0.99 |
| General Factor of Intelligence | 4291 | 0.03 | 1.01 | 2504 | 0.08 | 0.99 |
| Glucose (mmol/L) | 4297 | 4.78 | 0.58 | 2511 | 4.72 | 0.60 |
| HDL Cholesterol (mmol/L) | 4396 | 1.46 | 0.40 | 2546 | 1.48 | 0.41 |
| Neuroticism | 4426 | 3.60 | 3.09 | 2565 | 4.24 | 3.31 |
| Pack Years | 4380 | 7.54 | 14.30 | 2522 | 8.65 | 15.50 |
| QT Interval (corrected for heart rate; milliseconds) | 4374 | 7.70 | 24.20 | 2542 | 0.17 | 23.60 |
| Scottish Index of Multiple Deprivation (rank 1 : 6,505 = most to least deprived) | 4236 | 3980.00 | 1840.00 | 2457 | 3900.00 | 1890.00 |
| Systolic Pressure (mmHg) | 4447 | 134.00 | 18.40 | 2573 | 132.00 | 17.10 |
| Total Cholesterol (mmol/L) | 4409 | 5.21 | 1.05 | 2551 | 5.19 | 1.06 |
| Waist : Hip Ratio | 4383 | 0.87 | 0.09 | 2535 | 0.87 | 0.10 |

| *Ordinal Variables* | *n* | *Median* | *IQR* | *n* | *Median* | *IQR* |
|---|---|---|---|---|---|---|
| Educational Attainment (years) | 4303 | 4 | 3 | 2434 | 4 | 3 |
| *Self-Report Diseases* | *n* | *No. of Cases* | *% Cases* | *n* | *No. of Cases* | *%Cases* |
| Alzheimer's Disease Maternal | 4450 | 228 | 5.12% | 2531 | 128 | 5.06% |
| Alzheimer's Disease Paternal | 4450 | 134 | 3.01% | 2531 | 87 | 3.44% |
| Back Pain | 1693 | 480 | 28.35% | 1475 | 295 | 20.00% |
| Bowel Cancer | 4450 | 20 | 0.45% | 2531 | 11 | 0.43% |
| Breast Cancer | 4450 | 63 | 1.42% | 2531 | 40 | 1.58% |
| Chronic Kidney Disease | 4427 | 85 | 1.92% | 2563 | 40 | 1.56% |
| COPD | 4450 | 48 | 1.08% | 2531 | 32 | 1.26% |
| Depression | 4450 | 371 | 8.34% | 2531 | 414 | 16.36% |
| Diabetes | 4450 | 147 | 3.30% | 2531 | 89 | 3.52% |
| Heart Disease | 4450 | 196 | 4.40% | 2531 | 95 | 3.75% |
| Lung Cancer | 4450 | 5 | 0.11% | 2531 | 4 | 0.16% |
| Neck Pain | 1693 | 431 | 25.46% | 1475 | 257 | 17.42% |
| SCID Depression | 4450 | 825 | 18.54% | 2578 | 984 | 38.17% |
| Stroke | 4450 | 59 | 1.33% | 2531 | 41 | 1.62% |
