## Supplementary File 2 for "Epigenetic clocks predict prevalence and incidence of leading causes of death and disease burden"

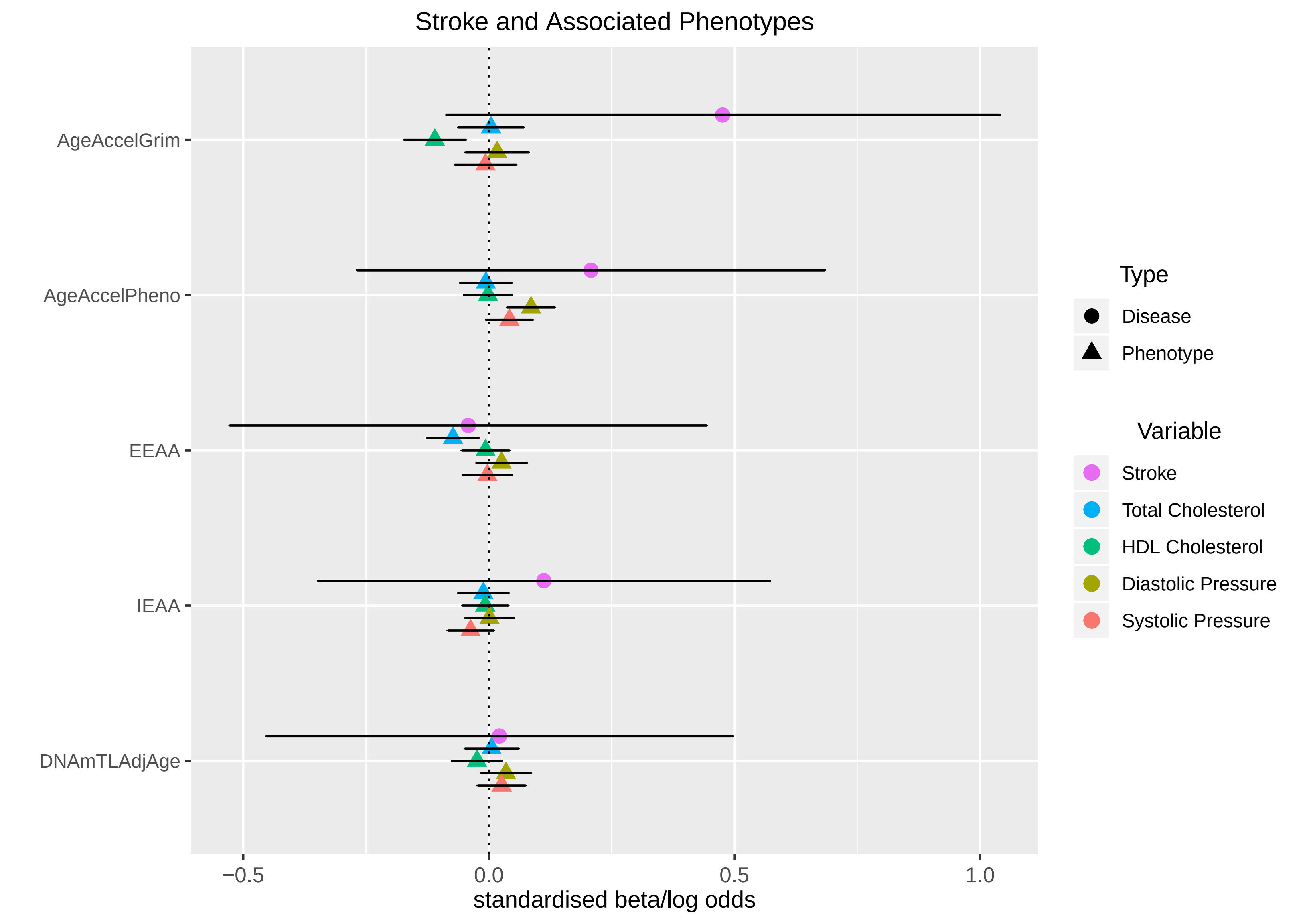
Stroke and Associated Phenotypes
AgeAccelGrim
AgeAccelPheno
EEAA
IEAA
DNAmTLAdjAge
−0.5
0.0
0.5
1.0
standardised beta/log odds
Type
Disease
Phenotype
Variable
Stroke
Total Cholesterol
HDL Cholesterol
Diastolic Pressure
Systolic Pressure

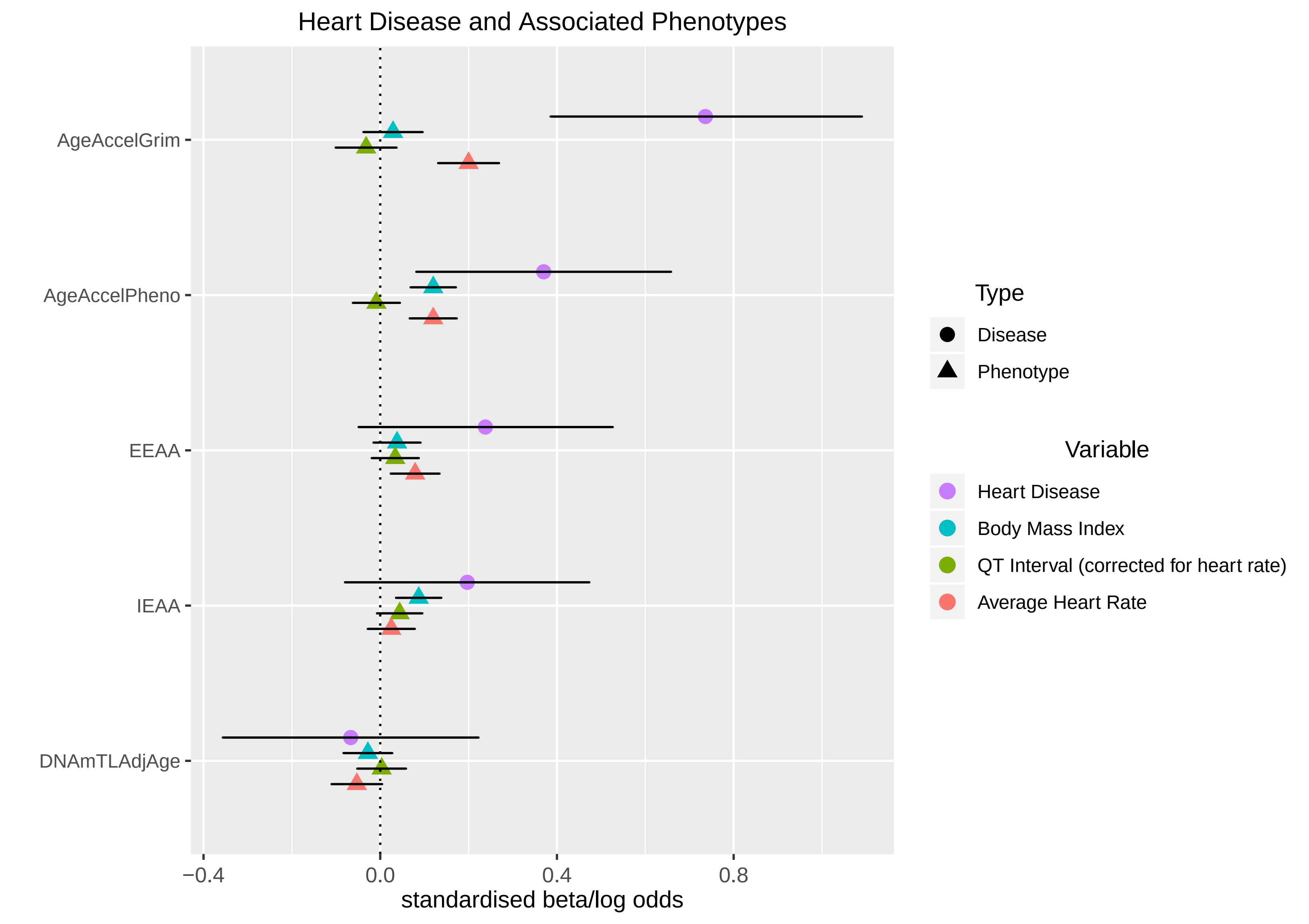

Heart Disease and Associated Phenotypes
AgeAccelGrim
AgeAccelPheno
EEAA
IEAA
DNAmTLAdjAge
−0.4
0.0
0.4
0.8
standardised beta/log odds
Type
Disease
Phenotype
Variable
Heart Disease
Body Mass Index
QT Interval (corrected for heart rate)
Average Heart Rate

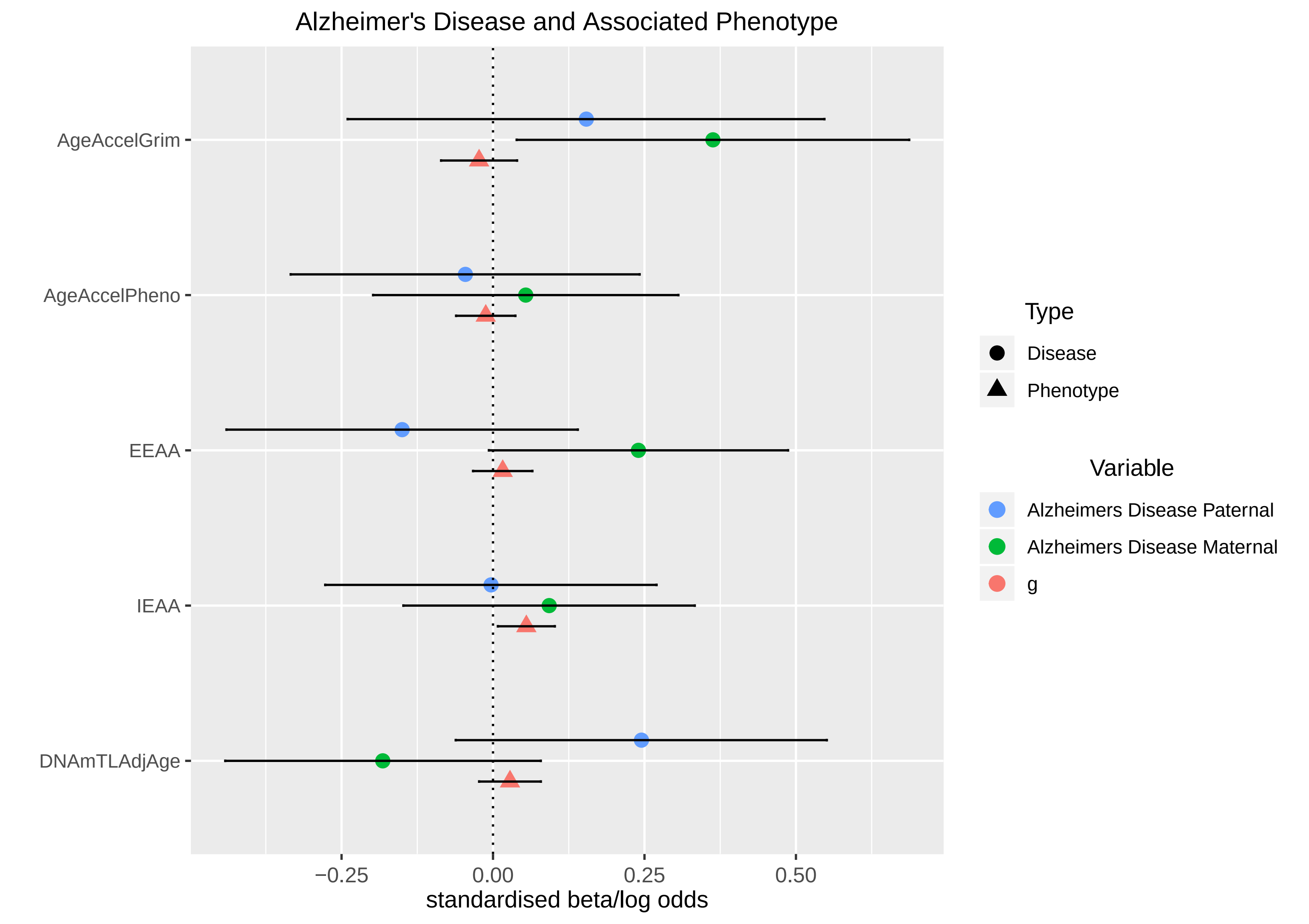

Alzheimer's Disease and Associated Phenotype
AgeAccelGrim
AgeAccelPheno
EEAA
IEAA
DNAmTLAdjAge
−0.25
0.00
0.25
0.50
standardised beta/log odds
Type
Disease
Phenotype
Variable
Alzheimers Disease Paternal
Alzheimers Disease Maternal
g

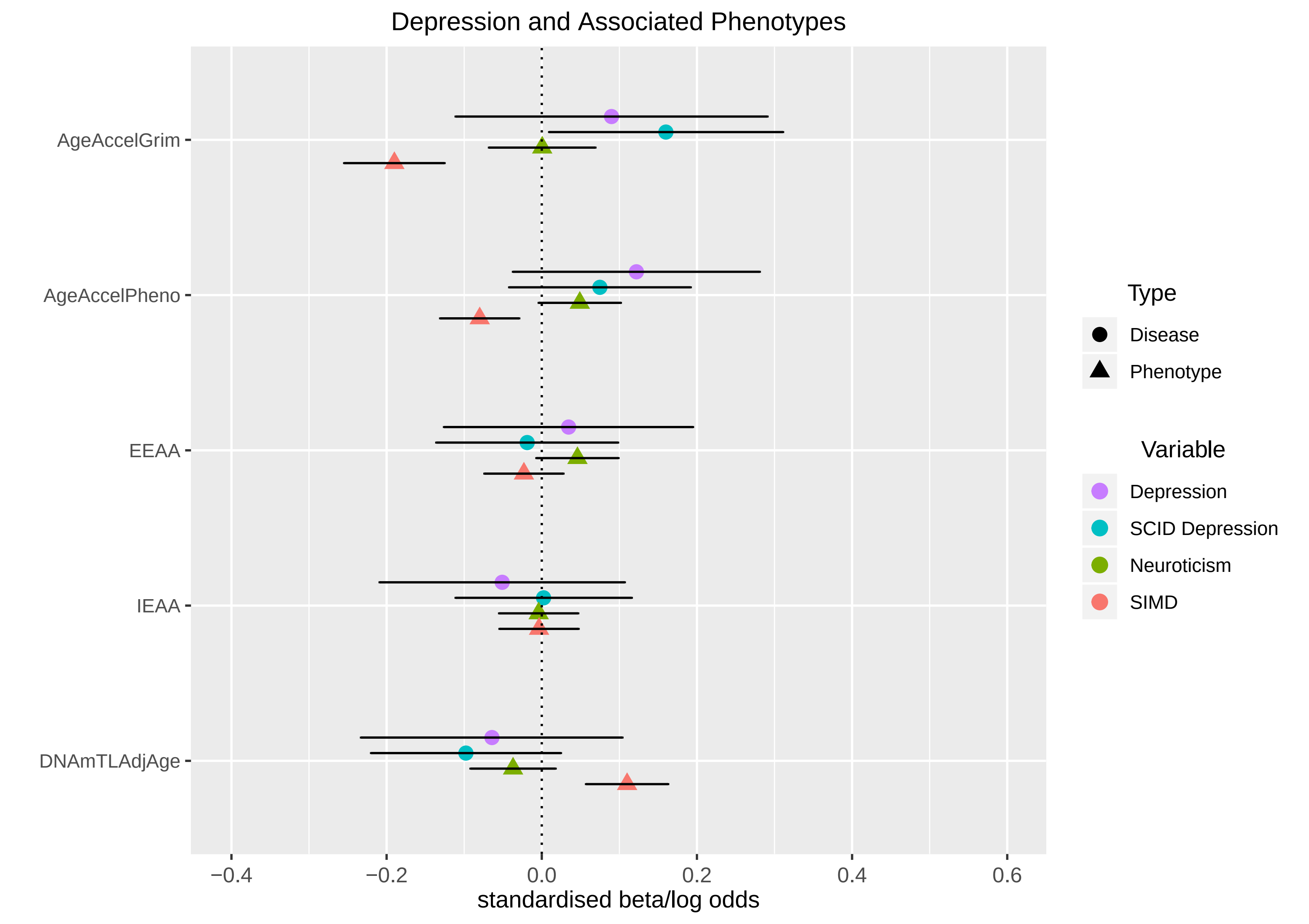
Depression and Associated Phenotypes
AgeAccelGrim
AgeAccelPheno
EEAA
IEAA
DNAmTLAdjAge
−0.4
−0.2
0.0
0.2
0.4
0.6
standardised beta/log odds
Type
Disease
Phenotype
Variable
Depression
SCID Depression
Neuroticism
SIMD

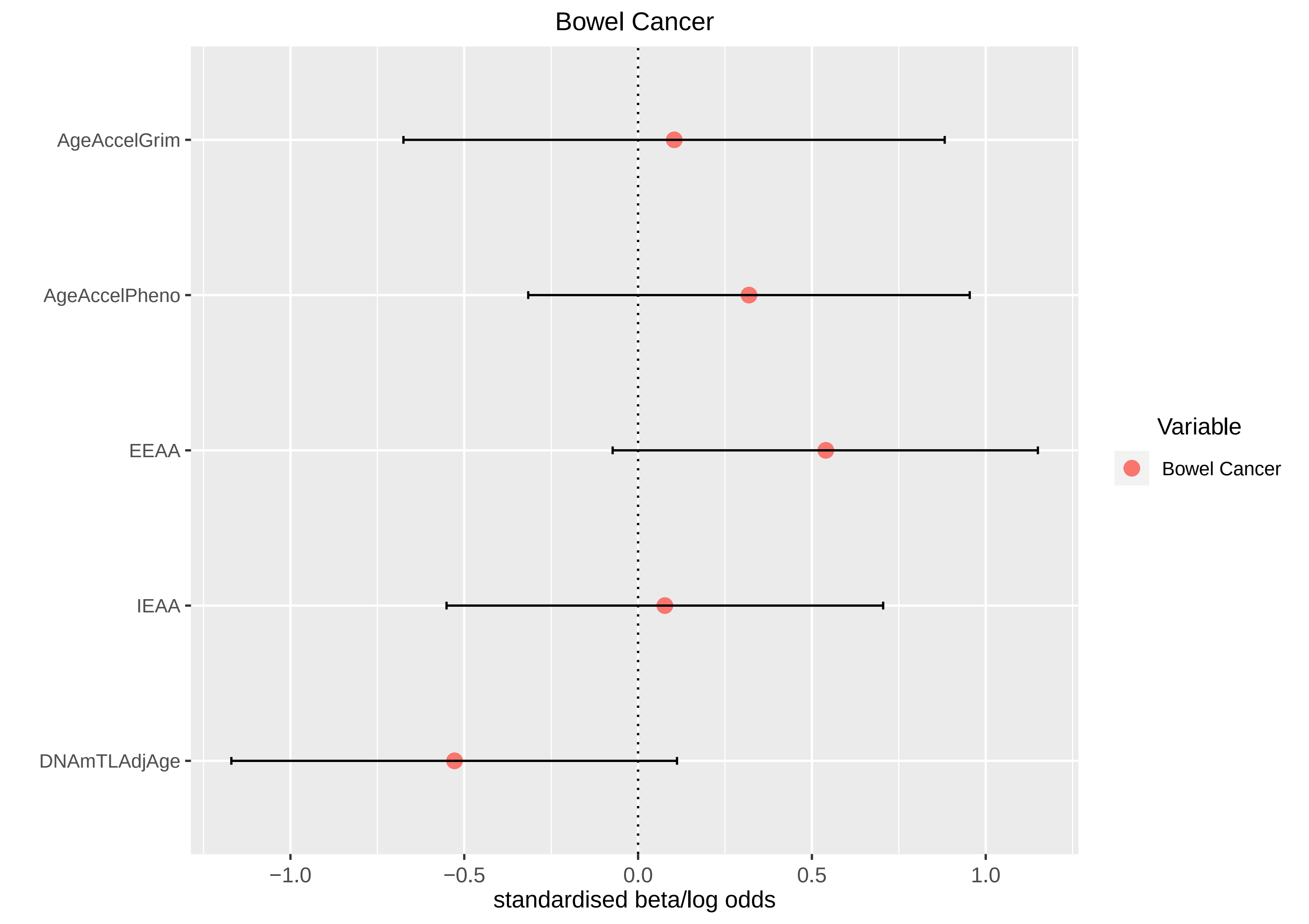

Bowel Cancer
AgeAccelGrim
AgeAccelPheno
EEAA
IEAA
DNAmTLAdjAge
−1.0
−0.5
0.0
0.5
1.0
standardised beta/log odds
Variable
Bowel Cancer

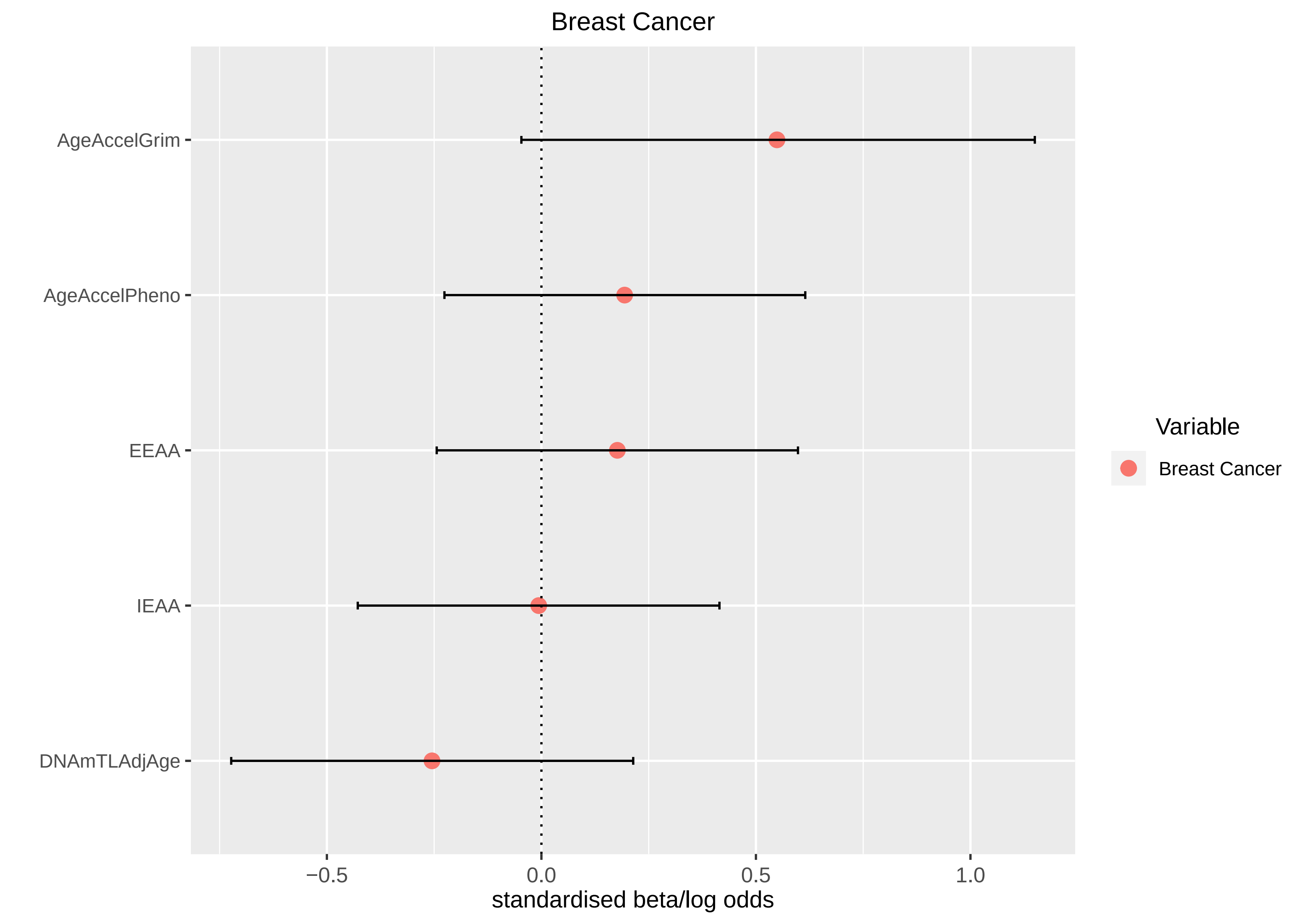
Breast Cancer
AgeAccelGrim
AgeAccelPheno
EEAA
IEAA
DNAmTLAdjAge
−0.5
0.0
0.5
1.0
standardised beta/log odds
Variable
Breast Cancer

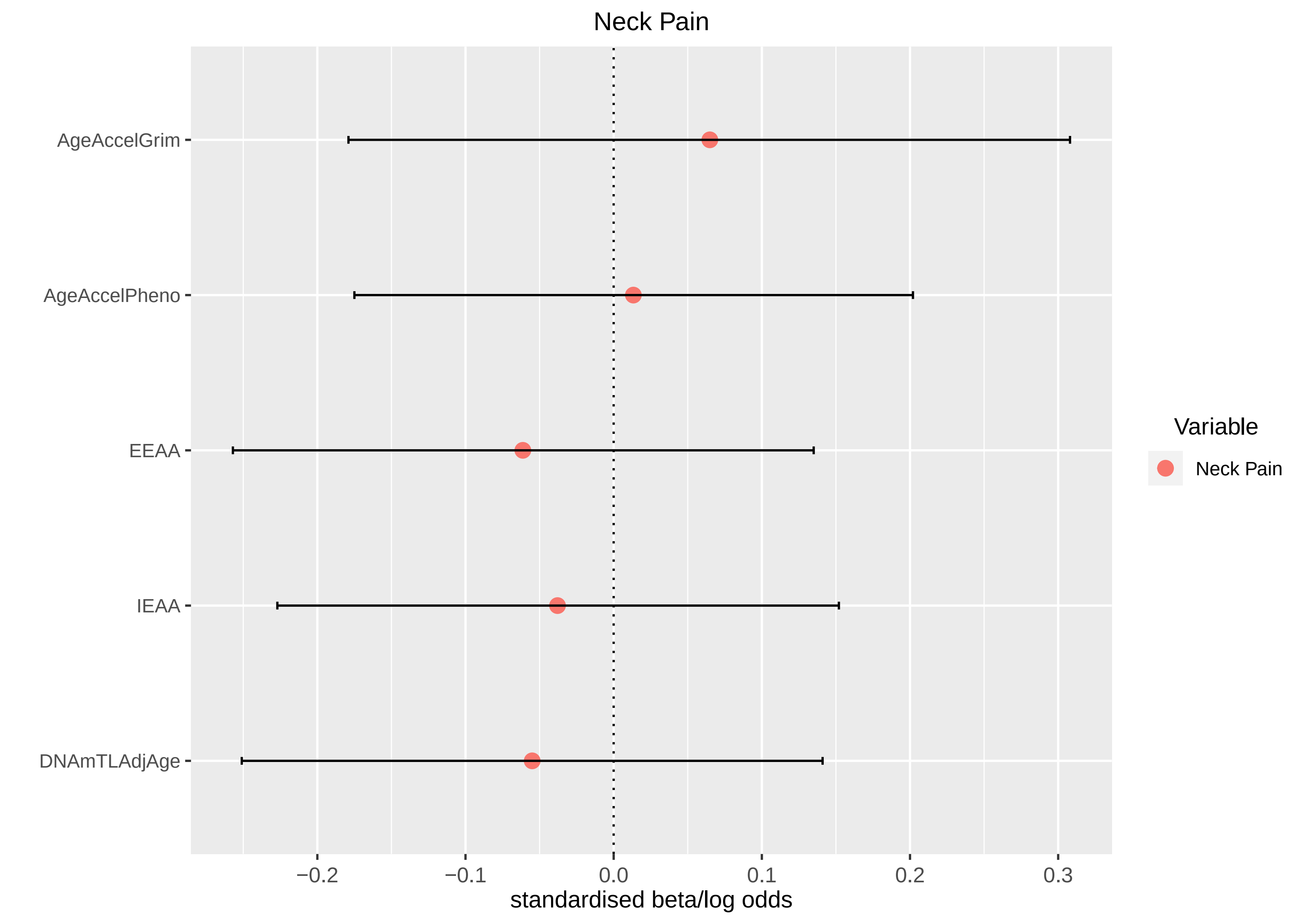
Neck Pain
AgeAccelGrim
AgeAccelPheno
EEAA
IEAA
DNAmTLAdjAge
−0.2
−0.1
0.0
0.1
0.2
0.3
standardised beta/log odds
Variable
Neck Pain

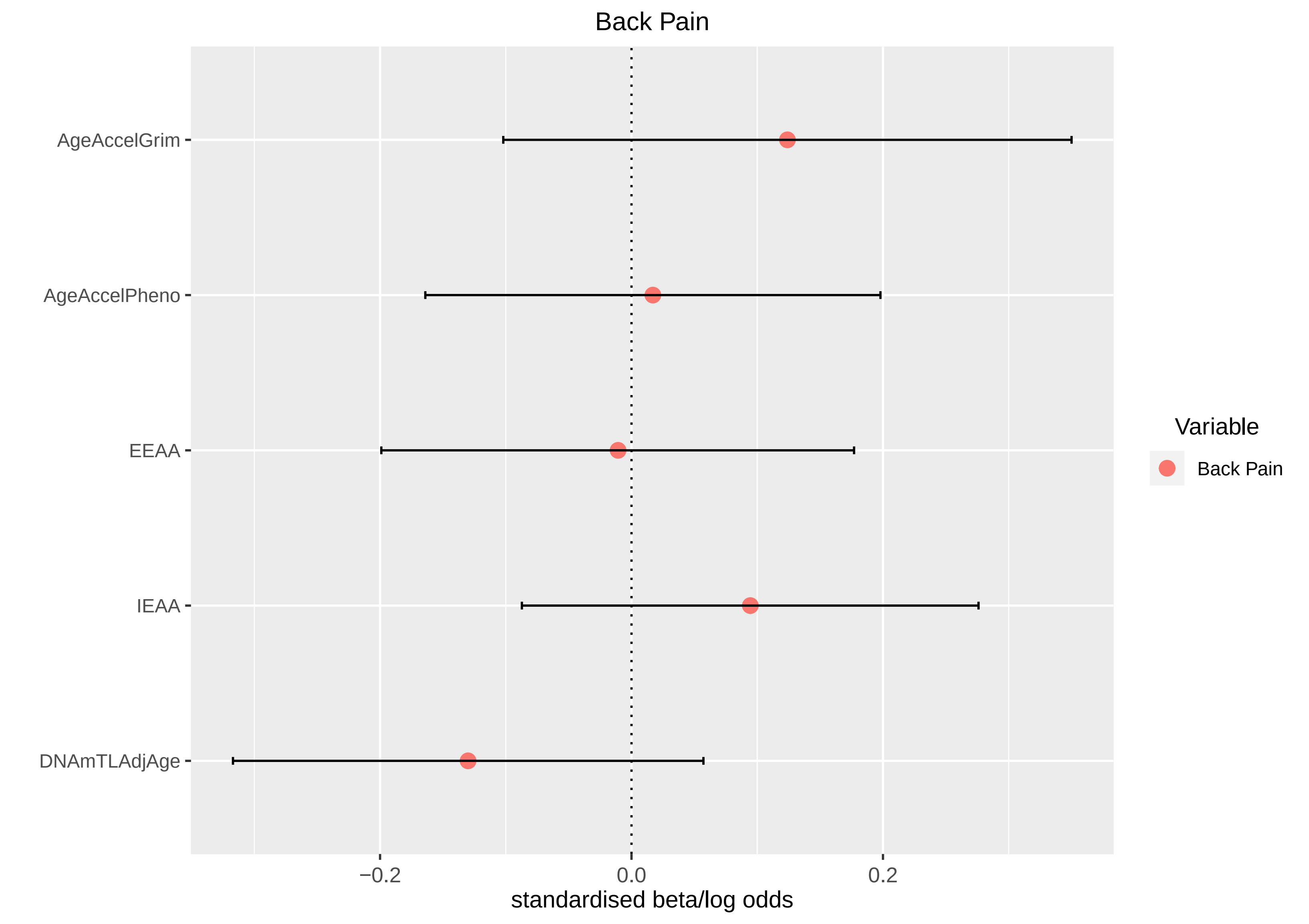
Back Pain
AgeAccelGrim
AgeAccelPheno
EEAA
IEAA
DNAmTLAdjAge
−0.2
0.0
0.2
standardised beta/log odds
Variable
Back Pain

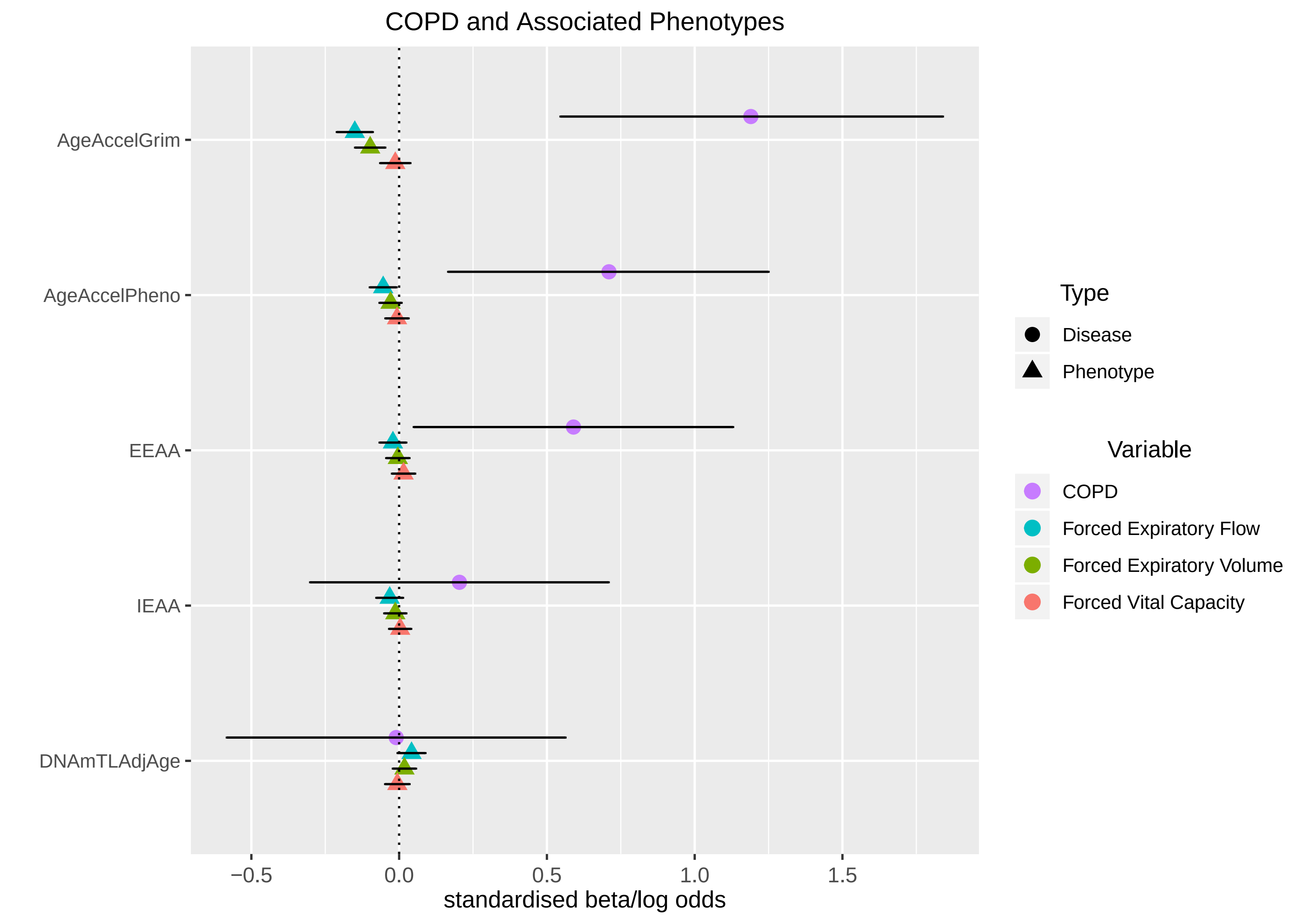
COPD and Associated Phenotypes
AgeAccelGrim
AgeAccelPheno
EEAA
IEAA
DNAmTLAdjAge
−0.5
0.0
0.5
1.0
1.5
standardised beta/log odds
Type
Disease
Phenotype
Variable
COPD
Forced Expiratory Flow
Forced Expiratory Volume
Forced Vital Capacity

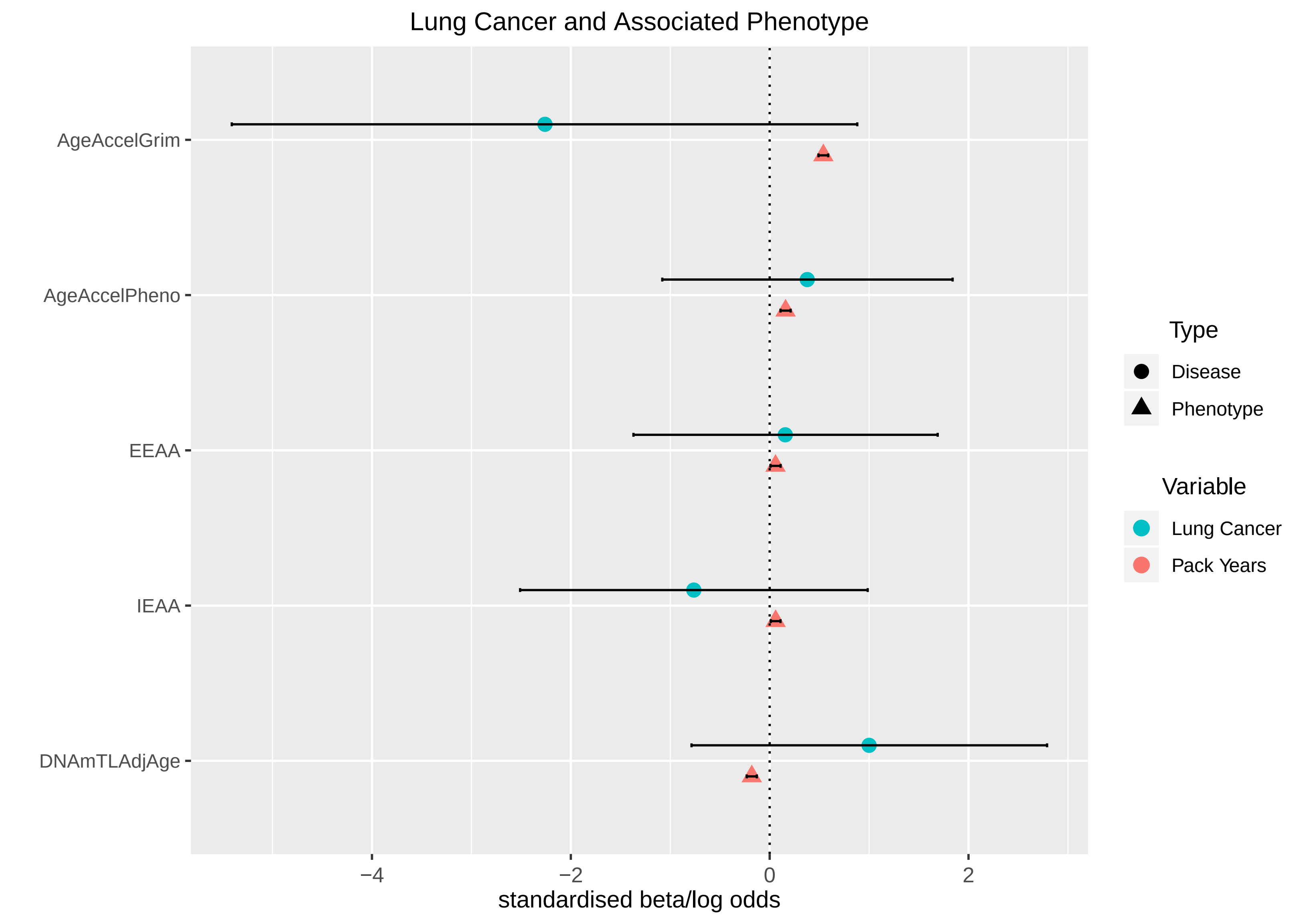
Lung Cancer and Associated Phenotype
AgeAccelGrim
AgeAccelPheno
EEAA
IEAA
DNAmTLAdjAge
−4
−2
0
2
standardised beta/log odds
Type
Disease
Phenotype
Variable
Lung Cancer
Pack Years

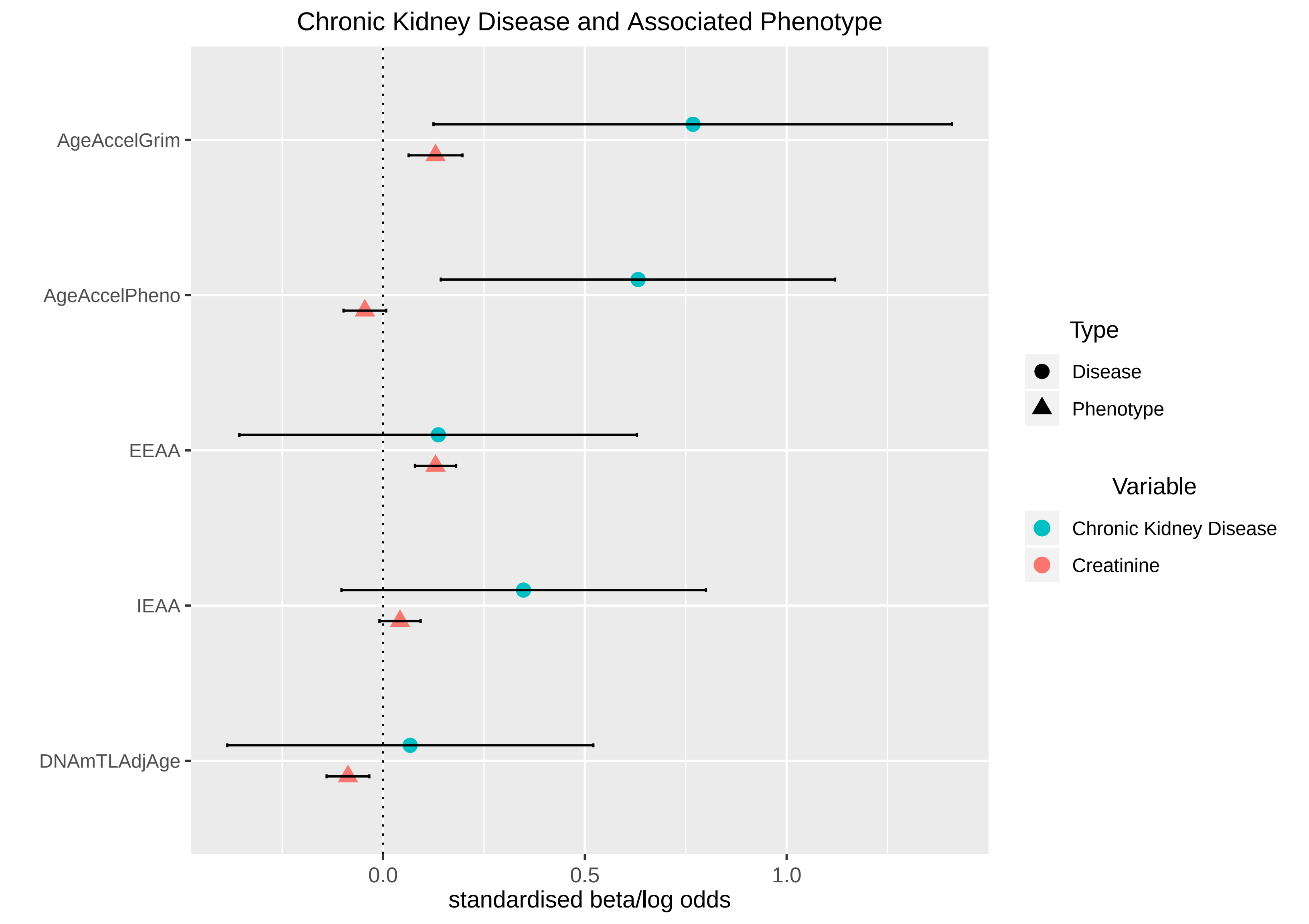
Chronic Kidney Disease and Associated Phenotype
AgeAccelGrim
AgeAccelPheno
EEAA
IEAA
DNAmTLAdjAge
0.0
0.5
1.0
standardised beta/log odds
Type
Disease
Phenotype
Variable
Chronic Kidney Disease
Creatinine

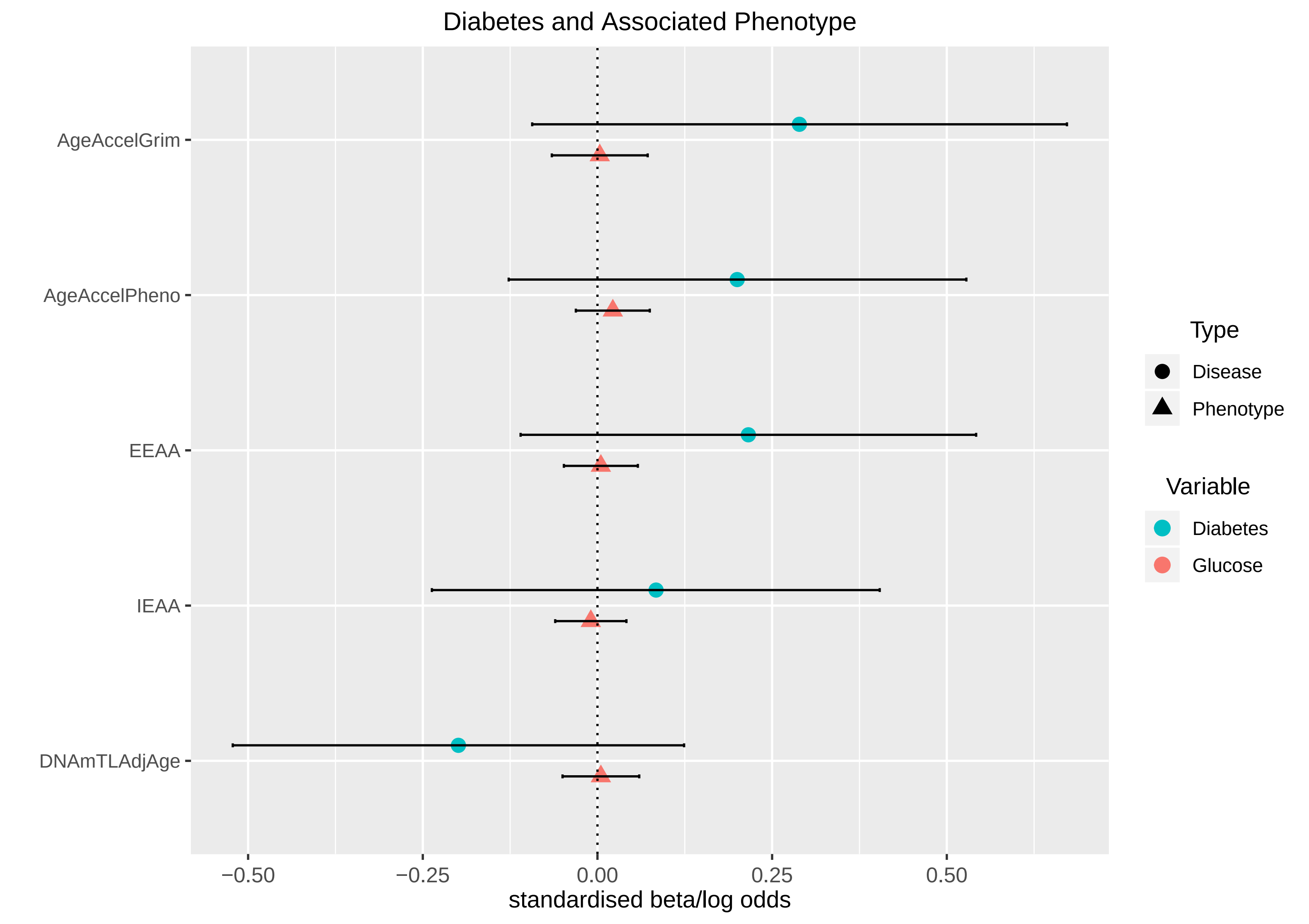
Diabetes and Associated Phenotype
AgeAccelGrim
AgeAccelPheno
EEAA
IEAA
DNAmTLAdjAge
−0.50
−0.25
0.00
0.25
0.50
standardised beta/log odds
Type
Disease
Phenotype
Variable
Diabetes
Glucose
