## Supplementary Methods for "Epigenetic clocks predict prevalence and incidence of leading causes of death and disease burden"

DNA methylation (DNAm) levels were measured using the Illumina HumanMethylationEPIC BeadChip Array on blood samples from Generation Scotland participants. Outliers were identified and excluded through visually inspecting a plot of log median intensity of methylated versus unmethylated signal. Samples were excluded in instances where ≥1% of CpG sites had a detection p-value in excess of 0.05, and where recorded sex did not match DNAm-predicted sex. Probes were removed in instances where ≥ 0.5% of samples had a detection p-value in excess of 0.05, and where there were beadcounts of less than 3 in 6% or more samples.

To derive measures of epigenetic age, we applied an additional quality control approach to those samples that had passed the screening process as described above. This was carried out to eliminate missing CpG values, as recommended in Horvath’s epigenetic clock tutorial (https://dnamage.genetics.ucla.edu/). Raw methylation IDAT files were read into *R* using the *minfi* package. Data were normalised through the normal-exponential convolution using out-of-band probe (noob) method implemented by the *preprocessNoob*() function in *minfi*. Briefly, this method consists of a background-subtraction step and dye-bias normalisation. Background noise is estimated from out-of-band probes and is removed from each sample whereas dye-bias normalisation utilises a subset of control probes to estimate the dye-bias. Noob-normalised methylation beta values were then obtained using the *getBeta*() function in *minfi*.

***Horvath Age***

Horvath Age gives rise to a measure of biological ageing termed “intrinsic epigenetic age acceleration (IEAA)” as it is independent of age-related changes in blood composition (1). An elastic net regression model was used to regress chronological age onto 21,369 probes that were present on both the Illumina 450k and 27k platforms and possessed fewer than 10 missing values. Penalised regression selected 353 CpGs as informative for predicting chronological age. The linear combination of these CpG sites provides an estimate of DNAm Horvath Age. Subsequently, IEAA was obtained as the residual term from regressing Horvath Age onto chronological age and fitting counts of naive CD8^+^ T-cells, exhausted CD8^+^ T-cells, plasmablasts, CD4^+^ T-cells, natural killer cells, monocytes, and granulocytes estimated from the methylation data.

***Hannum Age***

Hannum age produces a measure of epigenetic ageing referred to as “extrinsic epigenetic age acceleration (EEAA)” as it tracks age-related changes in blood cell composition (2). Elastic net regression identified 71 CpGs from the Illumina 450K platform as informative for predicting chronological age. EEAA was calculated in a two-step process. Firstly, a weighted average was calculated from Hannum Age and three cell types whose abundance exhibits age-related changes (naive cytotoxic T-cells, exhausted cytotoxic T-cells, and plasmablasts) (3). Secondly, EEAA was obtained by regressing this weighted average onto chronological age.

***DNAm PhenoAge***

DNAm PhenoAge was developed in a two-stage process by Levine *et al* (4). In the first step, a novel indicator of biological age, termed ‘Phenotypic Age’ was derived from the employment of an elastic net regression model in which the hazard of mortality was regressed onto 42 markers from the third National Health and Nutrition Examination Survey. In addition to chronological age, the model selected 9 haematological and biochemical markers for inclusion in the ‘Phenotypic Age’ predictor. These markers were: albumin, alkaline phosphatase, creatinine, C-reactive protein (CRP), mean cell volume, percentage lymphocytes, red cell distribution width, serum glucose (as indexed by glycated HbA1c) and white blood cell count. In the second stage, a second elastic net model was used to regress ‘Phenotypic Age’ onto 20,169 CpG sites (which represent those CpG sites present on all Illumina arrays i.e. 27K, 450K and EPIC). The model selected 513 CpG sites, the linear combination of which gives rise to a methylation-based surrogate of ‘Phenotypic Age’ termed ‘DNAm PhenoAge’. Importantly, the residual term from regressing DNAm PhenoAge onto chronological age produces a measure of biological age acceleration termed ‘AgeAccelPheno’.

**DNAm GrimAge**

In contrast to other clocks, which use chronological age as the dependent variable in elastic net regression models, Lu *et al.* developed a novel clock termed ‘DNAm GrimAge’ which uses mortality as a reference (5). They firstly created DNAm-based surrogates for smoking pack years and 88 plasma protein levels. DNAm-based surrogates were required to exhibit a moderate correlation coefficient of > 0.35 with the respective phenotype to be considered for stage two. Only 12 out of 88 proteins satisfied this condition. Consequently, for the second stage, an elastic net Cox regression model was used to regress time to death due to all-cause mortality onto chronological age, sex and methylation-based surrogates for smoking pack years and 12/88 proteins. The model selected chronological age, sex, DNAm smoking pack years and DNAm surrogates for 7/12 protein levels: adrenomedulin (DNAm ADM), beta-2-microglobulin (DNAm B2M), cystatin C (DNAm Cystatin C), growth differentiation factor 15 (DNAM GDF-15), leptin (DNAm leptin), plasminogen activation inhibitor 1 (DNAm PAI-1), and tissue inhibitor metalloproteinaise (DNAm TIMP-1). DNAm-based surrogates for smoking pack years and the 7 plasma proteins totalled 1,030 unique CpG sites from those present on both the Illumina 450K and EPIC array platforms. The linear combination of these variables provides an estimate of an individual’s DNAm GrimAge which correlates strongly with risk of mortality. The residual term from regressing DNAm GrimAge onto chronological age provides an index of biological age acceleration with respect to mortality risk and is termed ‘AgeAccelGrim’.

***DNAm Telomere Length***

Lu *et al.* developed an epigenetic correlate of telomere length in leukocytes, which is known to associate with a variety of age-related conditions. This biomarker termed ‘DNAm Telomere Length’ or ‘DNAm TL’ was derived using an elastic net regression model (6). In a similar design to DNAm GrimAge, CpG sites present on the Illumina 450K and Illumina EPIC Array platforms were regressed onto leukocyte telomere length. The model selected 140 CpG sites, and the weighted linear combination of methylation values at these CpG sites produced an individual’s estimate of DNAm TL. Interestingly, DNAm TL exhibited a higher correlation with age than measured leukocyte telomere length and outperformed its phenotypic counterpart in predicting time-to-death, time-to-coronary heart disease and time-to-congestive heart failure. A novel index of biological ageing was then derived by regressing DNAm TL onto chronological age which was termed ‘DNAmTLadjAge’. In contrast to other clocks, a lower age-adjusted DNAm TL is anticipated to correlate with poorer health outcomes as this reflects the age-related process of cellular telomere shortening.
