## Supplementary Note 1 for "Epigenetic clocks predict prevalence and incidence of leading causes of death and disease burden"

**Supplementary Note 1. The cross-sectional association of significant phenotypes with epigenetic clocks in discovery and replication cohorts in basic model adjusting for age and sex.**

*Cardiovascular Diseases*

AgeAccelGrim alone was associated with heart disease (Odds Ratio (OR) per SD = 1.71, P = 1.0 x 10^-7^) but not stroke following replication.

In relation to continuous traits, AgeAccelGrim, AgeAccelPheno and EEAA were positively associated with average heart rate (β = [0.08, 0.18], P = [1.1 x 10^-4^, 1.1 x 10^-19^]), with AgeAccelGrim showing the strongest association. AgeAccelPheno and IEAA were positively associated with body mass index (β = 0.11 and 0.09, P = 2.7 x 10^-8^ and 1.2 x 10^-5^, respectively). AgeAccelPheno was positively associated with higher diastolic blood pressure (β = 0.10, P = 2.4 x 10^-7^). AgeAccelGrim was negatively associated with high-density lipoprotein cholesterol (β = -0.11, P = 1.9 x 10^-8^). AgeAccelGrim and AgeAccelPheno were positively associated with waist-to-hip ratio (β = 0.15 and 0.10, P = 1.1 x 10^-16^ and 1.6 x 10^-8^, respectively). DNAmTLadjAge was negatively associated with average heart rate (β = -0.08, P = 1.7 x 10^-4^) and waist-to-hip ratio (β = -0.08, P = 7.6 x 10^-6^).

*Neurological and Psychiatric Diseases*

AgeAccelGrim was associated with both self-reported depression (OR = 1.26, P = 8.7 x 10^-5^) and SCID Depression (OR = 1.18, P = 6.6 x 10^-5^). No clock was associated with either maternal or paternal history of Alzheimer’s disease.

AgeAccelGrim was negatively associated with general cognitive ability (β = -0.14, P = 1.2 x 10^-11^) and a general factor of fluid intelligence (β = -0.14, P = 1.5 x 10^-12^). AgeAccelGrim alone was associated with neuroticism (β = 0.11, P = 4.8 x 10^-8^) in both cohorts. AgeAccelGrim and AgeAccelPheno were negatively associated with the Scottish Index of Multiple Deprivation (SIMD; lower indices correspond to more deprivation) (β = -0.27 and -0.12, P = 1.7 x 10^-38^ and 1.0 x 10^-9^, respectively). In contrast, DNAmTLadjAge was positively associated with SIMD (β = 0.15, P = 4.6 x 10^-13^).

*Pulmonary Diseases*

AgeAccelGrim alone was associated with COPD (OR = 3.49, P = 1.4 x 10^-13^). No clock was associated with lung cancer (no. of events = 5).

AgeAccelGrim and AgeAccelPheno were negatively associated with forced expiratory flow (β = -0.18 and -0.08, P = 2.6 x 10^-23^ and 1.7 x 10^-5^) and forced expiratory volume (β = -0.15 and -0.06, P = 5.7 x 10^-24^ and 4.3 x 10^-5^, respectively). AgeAccelGrim was also negatively associated with forced vital capacity (β = -0.08, P = 6.5 x 10^-8^). AgeAccelPheno was positively associated with smoking pack years (β = 0.17, P = 1.4 x 10^-17^). DNAmTLadjAge was positively associated with forced expiratory flow (β = 0.08, P = 1.4 x 10^-5^), forced expiratory volume (β = 0.07, P = 5.4 x 10^-6^) and negatively associated with pack years (β = -0.20, P = 2.5 x 10^-23^).

*Diabetes Mellitus and Kidney Disease*

AgeAccelGrim and AgeAccelPheno were associated with diabetes (OR: 1.47 and 1.48, P = 1.1 x 10^-4^ and 8.7 x 10^-5^, respectively).

AgeAccelGrim and EEAA were positively associated with creatinine (β = 0.08 and 0.14, P = 4.6 x 10^-5^ and 5.0 x 10^-13^, respectively). DNAmTLadjAge was negatively associated with creatinine (β = -0.11, P = 2.0 x 10^-8^).

*Cancer*

No clock was associated with either bowel or breast cancer following multiple testing correction.

*Neck and Back Pain*

AgeAccelGrim alone was associated with back pain (OR: 1.34, P = 5.4 x 10^-6^).

Table 1. Significant relationships of phenotypes with epigenetic clocks present in both discovery and replication cohorts in a basic model. Those phenotypes which remained significant in both cohorts in a subsequent fully-adjusted model are emboldened.

|  |  | Discovery Cohort | | | Replication Cohort | | |
| --- | --- | --- | --- | --- | --- | --- | --- |
| *Categorical Phenotypes* | | | | | | | |
| Clock | Variable | n event | OR | P | n event | OR | P |
| GrimAge | SCID Depression | 825 | 1.45 | 1.8 x 10^-22^ | 984 | 1.18 | 6.6 x 10^-05^ |
| GrimAge | Depression | 371 | 1.62 | 2.5 x 10^-22^ | 414 | 1.26 | 8.7 x 10^-05^ |
| **GrimAge** | **COPD** | **48** | **2.37** | **1.4 x 10^-12^** | **32** | **3.49** | **1.4 x 10^-13^** |
| GrimAge | Heart Disease | 196 | 1.54 | 5.2 x 10^-10^ | 95 | 1.71 | 1.0 x 10^-07^ |
| GrimAge | Diabetes | 147 | 1.58 | 1.4 x 10^-09^ | 89 | 1.47 | 1.1 x 10^-04^ |
| PhenoAge | Diabetes | 147 | 1.55 | 2.3 x 10^-08^ | 89 | 1.48 | 8.70 x 10^-05^ |
| GrimAge | Back Pain | 480 | 1.26 | 1.7 x 10^-05^ | 295 | 1.34 | 5.4 x 10^-06^ |
| *Continuous Phenotypes* | | | | | | | |
| Clock | Variable | n | β | P | n | β | P |
| **GrimAge** | **SIMD** | **4236** | **-0.28** | **3.3 x 10^-72^** | **2457** | **-0.27** | **1.7 x 10^-38^** |
| **HannumAge** | **Creatinine** | **4236** | **0.24** | **2.8 x 10^-62^** | **0.14** | **0.02** | **5.0 x 10^-13^** |
| **GrimAge** | **FEV** | **4427** | **-0.16** | **1.1 x 10^-42^** | **2191** | **-0.15** | **5.7 x 10^-24^** |
| GrimAge | g | 3776 | -0.22 | 6.1 x 10^-42^ | 2504 | -0.14 | 1.2 x 10^-11^ |
| **GrimAge** | **Average Heart Rate** | **4291** | **0.20** | **1.6 x 10^-37^** | **2572** | **0.18** | **1.1 x 10^-19^** |
| GrimAge | gf | 4444 | -0.19 | 5.9 x 10^-37^ | 2529 | -0.14 | 1.5 x 10^-12^ |
| GrimAge | Waist:Hip Ratio | 4324 | 0.15 | 6.5 x 10^-33^ | 2535 | 0.15 | 1.1 x 10^-16^ |
| **GrimAge** | **FEF** | **4383** | **-0.16** | **1.8 x 10^-29^** | **2185** | **-0.18** | **2.6 x 10^-23^** |
| **DNAmTL** | **Pack Years** | **3750** | **-0.17** | **3.0 x 10^-28^** | **2522** | **-0.20** | **2.5 x 10^-23^** |
| **PhenoAge** | **Body Mass Index** | **4380** | **0.16** | **1.4 x 10^-26^** | **2567** | **0.11** | **2.7 x 10^-08^** |
| **PhenoAge** | **Pack Years** | **4423** | **0.15** | **2.4 x 10^-24^** | **2522** | **0.17** | **1.4 x 10^-17^** |
| GrimAge | FVC | 4380 | -0.12 | 1.1 x 10^-22^ | 2191 | -0.08 | 6.5 x 10^-08^ |
| PhenoAge | Waist:Hip Ratio | 3775 | 0.11 | 1.3 x 10^-20^ | 2535 | 0.10 | 1.6 x 10^-08^ |
| **GrimAge** | **HDL Cholesterol** | **4383** | **-0.14** | **1.4 x 10^-20^** | **2543** | **-0.11** | **1.9 x 10^-08^** |
| **GrimAge** | **Creatinine** | **4396** | **0.14** | **2.5 x 10^-20^** | **2563** | **0.08** | **4.6 x 10^-05^** |
| **PhenoAge** | **Average Heart Rate** | **4427** | **0.13** | **3.3 x 10^-19^** | **2572** | **0.14** | **1.6 x 10^-13^** |
| DNAmTL | SIMD | 4444 | 0.13 | 5.5 x 10^-17^ | 2457 | 0.15 | 4.6 x 10^-13^ |
| DNAmTL | Creatinine | 4236 | -0.11 | 3.3 x 10^-14^ | 2563 | -0.11 | 2.0 x 10^-08^ |
| PhenoAge | SIMD | 4291 | -0.11 | 7.7 x 10^-14^ | 2457 | -0.12 | 1.0 x 10^-09^ |
| HannumAge | Average Heart Rate | 4236 | 0.11 | 9.2 x 10^-13^ | 2572 | 0.08 | 1.1 x 10^-04^ |
| PhenoAge | FEV | 4423 | -0.07 | 1.5 x 10^-09^ | 2191 | -0.06 | 4.3 x 10^-04^ |
| DNAmTL | Average Heart Rate | 3776 | -0.09 | 4.4 x 10^-09^ | 2572 | -0.08 | 1.7 x 10^-04^ |
| GrimAge | Neuroticism | 4324 | 0.08 | 5.7 x 10^-08^ | 2565 | 0.11 | 4.8 x 10^-08^ |
| DNAmTL | FEV | 4236 | 0.05 | 3.9 x 10^-06^ | 2191 | 0.06 | 5.4 x 10^-06^ |
| PhenoAge | Diastolic Pressure | 3776 | 0.06 | 4.3 x 10^-06^ | 2573 | 0.10 | 2.4 x 10^-07^ |
| DNAmTL | Waist:Hip Ratio | 4423 | -0.06 | 6.9 x 10^-06^ | 2535 | -0.08 | 7.6 x 10^-07^ |
| **HorvathAge** | **Body Mass Index** | **4383** | **0.06** | **4.2 x 10^-05^** | **2567** | **0.09** | **1.2 x 10^-05^** |
| PhenoAge | FEF | 4423 | -0.06 | 4.6 x 10^-05^ | 2185 | -0.07 | 1.7 x 10^-05^ |
| DNAmTL | FEF | 3750 | 0.06 | 5.0 x 10^-05^ | 2185 | 0.07 | 1.4 x 10^-04^ |
| *Mortality Analysis* | | | | | | | |
| Clock | Variable | n event | HR | P | n events | HR | P |
| GrimAge | **All-Cause Mortality** | **182** | **1.87** | **<2.0 x 10^-05^** | **57** | **1.7** | **6.5 x 10^-05^** |
| PhenoAge | All-Cause Mortality | 182 | 1.47 | 2.9 x 10^-09^ | 57 | 1.38 | 5.0 x 10^-03^ |
| DNAmTL | All-Cause Mortality | 182 | 0.70 | 6.4 x 10^-08^ | 57 | 0.81 | 0.01 |
| HannumAge | All-Cause Mortality | 182 | 1.39 | 1.1 x 10^-05^ | 57 | 1.39 | 3.0 x 10^-03^ |

COPD (chronic obstructive pulmonary disease), FEF (forced expiratory flow), FEV (forced expiratory volume), FVC (forced vital capacity), g (general factor of cognitive ability), gf (general factor of fluid intelligence), HDL (high-density lipoprotein), HR (hazard ratio), OR (odds ratio), SCID (Structured Clinical Interview for DSM), SIMD (Scottish Index of Multiple Deprivation).
