## Supplementary Note 2 for "Epigenetic clocks predict prevalence and incidence of leading causes of death and disease burden"

**Supplementary Note 2. The longitudinal association of ICD-10-coded common diseases with epigenetic clocks in a basic model adjusting for age and sex.**

*DNAm GrimAge*

AgeAccelGrim was associated with incidence of COPD (Hazard Ratio (HR) per SD = 2.70, P < 2.0 x 10^-16^), diabetes (HR = 1.67, P < 2.0 x 10^-16^), ischemic heart disease (HR = 1.48, P = 1.1 x 10^-14^), stroke (HR = 1.44, P = 4.1 x 10^-7^) and lung cancer (HR = 1.60, P = 1.5 x 10^-14^). AgeAccelGrim also showed a nominally significant association with incidence of depression (HR = 1.42, P = 3.3 x 10^-3^), Alzheimer’s disease (HR = 1.64, P = 0.01) and dorsalgia (HR = 1.23, P = 0.02).

*DNAm PhenoAge*

AgeAccelPheno was associated with incidence of diabetes (HR = 1.62, P = 7.7 x 10^-16^), COPD (HR = 1.55, P = 2.5 x 10^-10^) and ischemic heart disease (HR = 1.28, P = 1.4 x 10^-6^). AgeAccelPheno showed nominally significant associations with incidence of depression (HR = 1.45, P = 9.4 x 10^-3^) and lung cancer (HR = 1.15, P = 0.04).

*HannumAge*

Age-adjusted HannumAge (EEAA) was associated with incidence of COPD (HR = 1.27, P = 3.1 x 10^-4^).

*HorvathAge*

Age-adjusted HorvathAge (EEAA) showed only a nominally significant relationship with incidence of diabetes (HR = 1.18, P = 7.6 x 10^-3^).

*DNAmTLadjAge*

Age-adjusted DNAm Telomere Length was associated with incidence of COPD (HR = 0.63, P = 6.3 x 10^-11^), ischemic heart disease (HR = 0.81, P = 5.7 x 10^-5^), stroke (HR = 0.75, P = 7.9 x 10^-5^) and diabetes (HR = 0.79, P = 3.3 x 10^-4^), and also showed a nominally significant association with incidence of depression after 13 years since study baseline (HR = 0.72, P = 0.03).

Refer to Supplementary Table 9 for full details of association models.
